# *Esrrb* is a cell cycle dependent XEN priming factor balancing between pluripotency and differentiation

**DOI:** 10.1101/2020.08.03.234112

**Authors:** Sapir Herchcovici Levi, Sharon Feldman, Lee Arnon, Shlomtzion Lahav, Muhammad Awawdy, Adi Alajem, Danny Bavli, Xue Sun, Yosef Buganim, Oren Ram

## Abstract

Cell cycle and differentiation decisions are tightly linked; however, the underlying principles that drive these decisions are not fully understood. Here, we combined cell-cycle reporter system and single-cell RNA-seq profiling to study the transcriptomes of mouse embryonic stem cells (ESCs) in the context of cell cycle states and differentiation. By applying retinoic acid, a multi-linage differentiation assay, on G1 and G2/M pre-sorted ESCs, we show that only G2/M ESCs were capable of differentiating into extraembryonic endoderm cells (XENs), whereas cells in the G1 phase predominantly produce Epiblast Stem Cells. We identified ESRRB, a key pluripotency factor that is upregulated during G2/M phase, as a central driver of XEN differentiation. Furthermore, enhancer chromatin states based on WT and *Esrrb*-KO ESCs show association of ESRRB with XEN poised enhancers. Cells engineered to overexpress Esrrb during G1 allow ESCs to produce XENs, while ESRRB-KO ESCs lost their potential to differentiate into XEN. In addition, Embryonic bodies (EBs) are not affected by deletion of ESRRB but trigger apoptosis upon attempts to apply direct XEN differentiation. Taken together, this study reveals an important functional link between Esrrb and cell-cycle states during the exit from pluripotency. Finally, the experimental scheme of single cell RNA-seq in the context of cell cycle can be further expanded into other cellular systems to better understand differentiation decisions and cancer models.

## Introduction

Embryonic stem cells (ESCs) are pluripotent cells derived from the inner cell mass of the preimplantation blastocyst ^1^. These cells possess unique properties of self-renewal and the ability to give rise to all definitive structures of the fetus. The first cellular decision distinguishes between the epiblast, which produces the embryo body, and the hypoblast, which contributes to the extraembryonic endoderm cells (XEN) ^2,3^.

The transition from an uncommitted to a differentiated state requires rapid and global execution of specific gene programs including the gradual silencing of pluripotency genes and the activation of lineage-specific genes. External signals sensed by each cell drive its fate decisions. ESCs must coordinately alter their transcriptomes, chromatin architectures, and epigenetic landscapes throughout the differentiation process^4–6^.

The cell cycle is a critical process in the development of an organism, and it is closely linked to cell-fate decisions^7,8^. Cell-cycle consists of four distinct phases dedicated to the replication and transmission of genetic material to daughter cells; in S-phase and M-phase cells, chromosome replication and chromosome transmission occur, respectively. These key events are separated by gap phases, G1 and G2, that serve as regulatory windows to ensure that cell-cycle events occur at the correct time and in the right order^9,10^. The ESC cell-cycle structure is characterized by a short G1 phase and a high proportion of cells in S phase ^11^. This is associated with pluripotency factors that influence activities of cyclin-dependent protein kinases (CDKs) ^12^. Studies have shown that the characteristic cell-cycle of ESCs is also affected by the culture conditions. ESCs cultured in serum lack G1 control and rapidly progress through the cell cycle, whereas ESCs grown in the presence of two small molecule inhibitors (2i), MEK inhibitor (PD0325091) and Gsk3β inhibitor (CHIR99021), have a longer G1 phase ^13^.

Previous studies have shown that the cell-cycle stage is a major determinant of cell-fate decisions^8,14–19^. Evidence suggests that G1/S is the cellular stage at which differentiation decisions are made, whereas the G2 phase is mostly, but not only, dedicated to mitosis control ^20^. More specifically, Gonzales et. al. showed hESCs cell cycle dependent gradual response to differentiation ques. hESCs engage toward differentiation in early G1 and then decide only in G2 to fully commit toward a specific lineage by dissolving their pluripotent state ^21^. The intersection between cell-cycle regulation and cell-fate decision mechanisms involves developmental signals and CDK activities, which mediate cell-cycle dependent changes in the epigenetic landscape and chromosome architecture of developmental genes^22^. The CDKs are also responsible for recruitment of transcription factors to cell-fate related genes ^22^. Pauklin et al. showed that cell cycle regulators (as Cyclin D1–3) control cell fate decisions in human pluripotent stem cells by engaging transcriptional complexes onto different developmental genes ^23^.Transcription factor activities can counter CDK activities and drive the cells to exit the pluripotent state. Thus, the balance between CDK activity and transcription factor activity determines cell fate.

Retinoic acid (RA), the active metabolite of vitamin A, is crucial in early embryonic development and in maintenance of many organ systems in the adult organism ^24^. It has been shown that RA represses pluripotency-associated genes and activates lineage-specific markers in ESCs ^25^. RA promotes a variety of lineage outcomes such as ectoderma ^26–28^, endodermal and extraembryonic endoderm (XEN) cells ^29–33^. The ability of RA to promote various differentiation phenotypes implies that RA is involved in the switch between proliferation and differentiation^11,28^. Semrau, S. et al.^29^ further explored RA based exit from pluripotency at a single cell level. In their study they show that the exit from pluripotency can be traced by single cell RNA-seq after 24 hours with RA. After 96 hours, Ectodermal and XEN subpopulations arose. However, the link between ESCs cell cycle states, exit from pluripotency and the effect on differentiation outcomes has not been tested.

Pluripotent cell identity is sustained by the activity of a highly interconnected network of transcription factors such as *POU5F1, NANOG*, and *SOX2* ^34^ and a large group of ancillary factors such as Estrogen related receptor beta (Esrrb) ^35^. *Esrrb* is an orphan nuclear receptor that is required for self-renewal and pluripotency of ESCs ^36^. In ESCs, *Esrrb* function is controlled by extrinsic cues mediated by kinases such as GSK3i ^37^ and intrinsic regulators such as *Nanog* ^38^. This confers flexibility to the pluripotency network, as changes in the activity of these factors modulate the balance between maintenance and loss of pluripotency ^39^. In the early post-implantation mouse embryo, *Esrrb* is specifically expressed in the extraembryonic ectoderm and plays a crucial role in trophoblast development ^35^. Furthermore, Benchetrit et al ^40^ demonstrated that induced extraembryonic endoderm stem cells (iXENs) require high levels of *Esrrb*, pointing out the possible role of *Esrrb* in regulating XEN lineage differentiation.

Here, by combining a sensitive cell-cycle reporter system with scRNA-seq, we studied the cross-talk between cell-cycle state and cell-fate decisions during the exit from pluripotency. We revealed that following RA, ESCs in G1 exclusively differentiate to EpiSCs and mesodermal cells. Strikingly, if cells are exposed to RA during G2/M phase, they have the unique capacity to differentiate into primitive endoderm and more specifically into XENs. We identified that this capacity is driven by upregulation of *Esrrb* during G2/M state. Overexpressing Esrrb in G1 ESCs cells, enabled XEN differentiation. ESRRB-KO ESCs lost their potential to produce XEN cells. We show that this loss involves changes in ESCs enhancer landscape and more specifically the loss of H3K4me2 on endodermal enhancers. In addition, ESRRB ChIP-seq published by Festuccia, N. *et al* ^41^, further support the direct association of ESRRB with XEN enhancers. Overall, our results demonstrate that exit from pluripotency happen earlier during pluripotency, suggesting that cell cycle dependent pluripotency factors regulate cell fate decisions outcomes.

## Results

### ESCs differentiation is coupled with a decrease in proliferation and changes in cell-cycle state composition

In order to visualize and effectively distinguish between cell cycle states without complications of cell-to-cell variation, we produced a clonal ESC cell line expressing the FUCCI cell-cycle reporter system ^42^. The FUCCI system is based on cycled translation and ubiquitination-based degradation of Cdt1 (conjugated to mKO2, red) and Geminin (conjugated to mAG1, green). Nuclei of G1 cells appear orange, and there is a gradual increase in the intensity of green as cell-cycle progresses, with a peak at G2/M (Fig. 1A and B). The insertion of the lentiviral-based FUCCI probes into ESCs did not result in detectable differences in pluripotency or in the ability of cells to differentiate. To induce differentiation, we grew ESCs in RA-containing media for 4 days. We chose the RA differentiation protocol for its broad effects on differentiation outcomes, driving ESCs to differentiate towards intra-embryo progenitor cells (e.g. EpiSCs) but also towards extra-embryo cells such as XEN ^28^.

**Figure 1:**
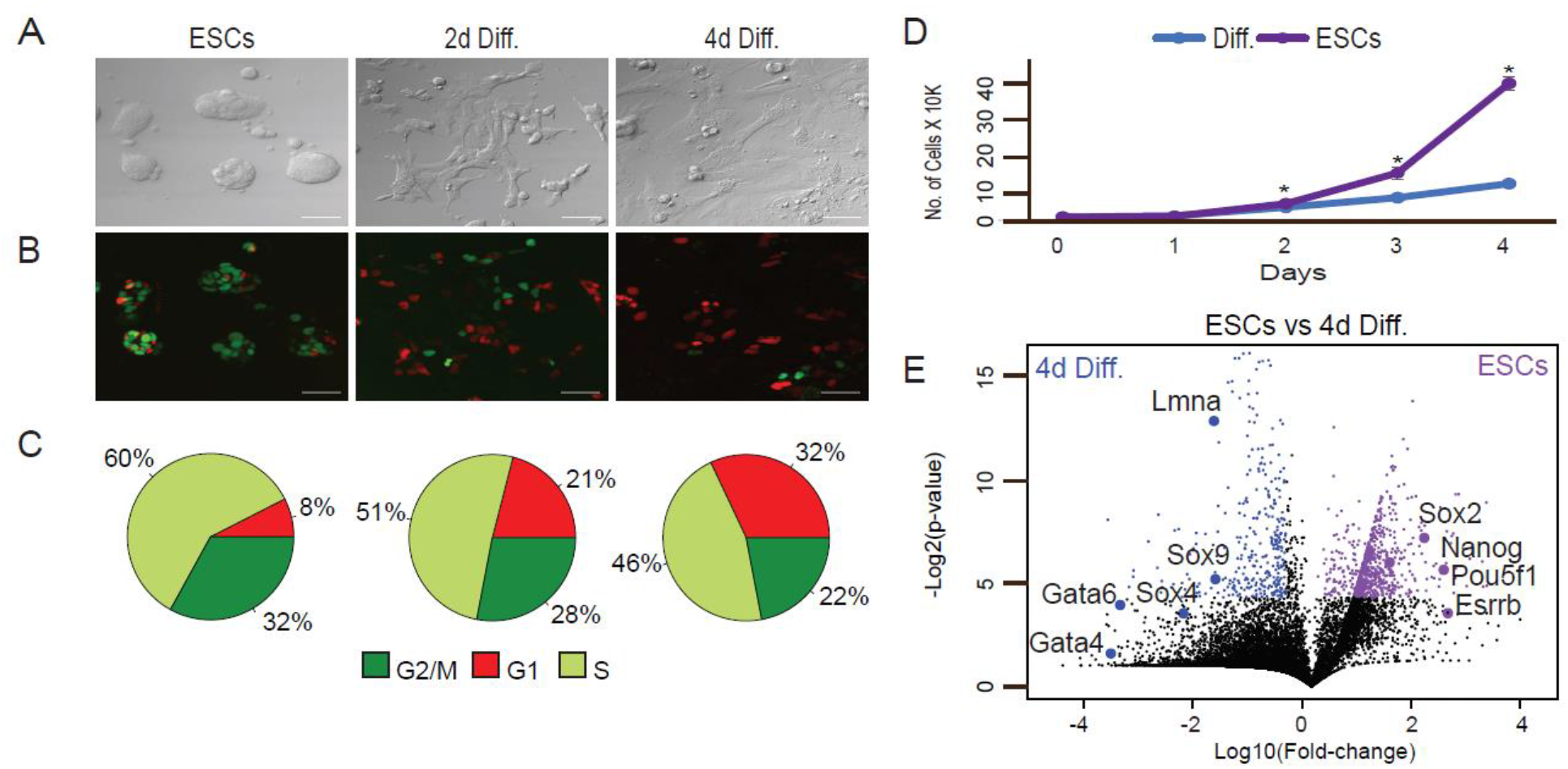
Temporal ESC FUCCI based system. ESCs were maintained in LIF/2i ESC medium and then switched to differentiation medium (1 µM RA without LIF and 2i) and analyzed after 2 and 4 days (2d Diff. and 4d Diff., respectively). **(A**) Representative X20 bright-field images, scale bar 50 µm. **(B**) Representative X20 confocal images, scale bar 50 µm. (**C**) Cell-cycle phases in pluripotent state and during differentiation calculated by a combination of FUCCI and Hoechst DNA staining. (**D**) Growth rates of ESCs and RA treated cells (diff.). Shown are mean values of two independent experiments (* p < 0.05, Wilcoxon test). (**E**) Differential expression of transcriptomes using RNA-seq. Transcriptomes of ESCs (WT) vs. cells treated with RA for 4 days (differentiated) were plotted. The x axis represents expression fold changes in logarithmic scales and the y axis represents p-values based on three replicates. Genes upregulated in ESCs are indicated in purple and genes upregulated in the differentiated cells are indicated in blue.

Upon RA treatment, cells underwent morphology changes typical of differentiation ^43^ (Fig. 1A and B). To ensure accurate separation between S and G2/M cellular states, we set the FACS sorting parameters using integration of both FUCCI and Hoechst staining of DNA (Fig. S1A). Confocal imaging and FACS analysis of the FUCCI reporters and Hoechst staining of DNA revealed that prior to addition of RA, approximately 8% of the cells were in the G1 phase; 60% in S and 32% in G2/M (Fig. 1C). As cells marched into differentiation, G1 cell proportion increased. After 2 and 4 days of RA treatment, 21% and 32% of the cells were captured in G1 phase, respectively. G1 increase is accompanied by replication rate reduction (Fig. 1D). As ESCs differentiate, cells pause in G1 phase, thus fewer replicating cells are captured ^11^. Proliferation rate analysis by CellTrace verified that the rate of cell divisions was significantly slower in cells grown in differentiation conditions than in pluripotent ESCs (Fig. S1B).

Over the 4-day period, we observed transcriptional activation of genes that regulate EpiSC- and XEN-like states in the embryo, such as *Sox4, Sox9, Lmna, Gata4*, and *Gata6*. Concomitantly, there was downregulation of pluripotency factors such as *Pou5f1, Nanog*, and *Esrrb* (Fig. 1E). Additional genes associated with differentiation or pluripotent state with altered expression over the time course are listed in Supplementary Table 1.

### XEN differentiation potential is driven by the cell-cycle phase of ESCs

To map the potential links between cell-cycle states during pluripotency and differentiation trajectories, ESCs were sorted into G1, and G2/M states. Immediately after sorting, we initiated differentiation by replacing the standard medium with medium containing RA (Fig. 2A). To analyze the heterogeneous population of cells resulting from RA treatment, we used the InDrop scRNA-seq system ^44^ and Seurat pipeline for analysis ^45^. Based on two biological replicates and analysis of transcriptomes of ∼2500 cells, we detected four main subpopulations (Fig. 2B): (1) XEN cells (cluster 0), (2) EpiSCs (clusters 1 and 3), and (3) mesodermal progenitor cells (cluster 2). We further eliminated the possibility that cluster separation could be simply explained by cell coverage (Fig. 2C). Cluster 3 presented slightly higher coverage compared to the other clusters, however it shows overall similar markers as cluster 1, both supporting EpiSC differentiation states and thus not compromising our biological interpretations. Next, we revealed that G1 cells predominantly exhibit epiblast differentiation capacities, with expression profiles characteristic of EpiSCs and mesodermal progenitor cells (Fig. 2D and 2E). In Fig. 2F, we marked EpiSC marker genes highly expressed in clusters 1 and 3 (*Sox4, Sox9, Sox11, Brd2, Nrp1, Vimentin, Ecm1, Smad6*, and *Gata2*). *Gata2, Sin3b, Rhox6 and Rhox9* specifically mark mesodermal subpopulation in cluster 2 (Fig. 2F) ^46^. Like G1 ESCs, also G2/M ESCs enabled differentiation towards EpiSC, however, most of G2/M ESCs differentiated into XEN cells that expressed primitive endoderm markers such as *Gata4, Foxq1, Foxa2, Dab2, Lama1 and Lamc1* (Fig. 2F). All marker genes based on the four clusters are listed in Supplementary Table 1. Finally, to rule out the possibility that upon LIF/2i withdrawal, high levels of BMPs in the serum may have an impact on the differentiation propensities, we repeated the experiment described in Fig2 with ESCs growing in a fully chemically defined medium, and obtained similar results (Fig S2). This suggests that ESC culturing method is not affecting differentiation potential and outcomes.

**Figure 2:**
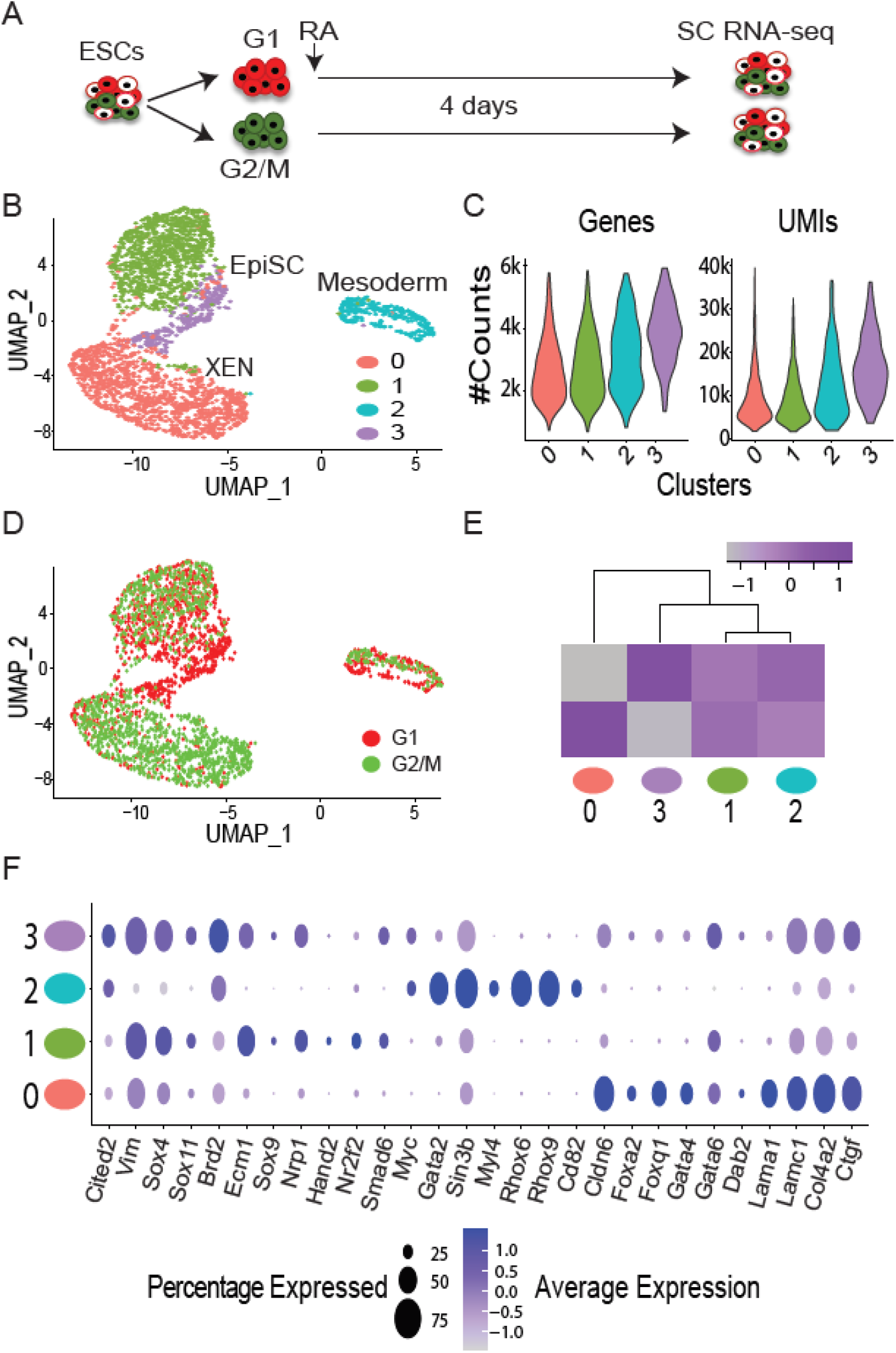
scRNA-seq analysis of differentiated cells based on G1 and G2/M ESCs populations. (**A**) Schematic illustration of the sorting and scRNA-seq experiments. (**B**) UMAP visualization of ∼2000 single cells clustered into four groups using the Seurat pipeline ^45^, EpiSC (marked as 1 and 3), XEN (marked as 0) and mesodermal-like cluster (marked as 2) are also represented by different colors. (**C**) Violin plots representing number of genes and UMIs for each cluster (**D**) UMAP visualization of the same single-cell data colored based on the initial sorting of ESCs; G1 are in red, and G2/M are in green. (**E**) Heatmap describing proportion of cells in each cluster (x axis) based on cell cycle-initiated ESC populations (y axis). Gray to purple scale indicates normalized cell number in each cluster. (**F**) Dot plot of differentially expressed genes explaining the four different clusters. Dot size indicates percentage of cells that express the gene and gray to purple scale indicates average expression within the cluster.

Next, we tried to capture the dynamics of RA based differentiation. We applied diffusion map to examine unsorted ESCs together with G1 vs G2/M sorted ESCs followed by 2 and 4 days of RA differentiation. Diffusion map analysis clearly shows that 2 days of differentiation support a very early step of differentiation compared to the major clusters observed after 4 days (Fig. S3). Interestingly, following 4 days of differentiation, endoderm and XEN states are spatially adjacent on diffusion map space, but we could still distinguish between these states, supporting the observation that only G2/M ESCs hold the capacity to produce XEN cells (Fig. S3D and S3G).

Overall, these results show that the differentiation capacity of ESCs is strongly influenced by the state of the cell-cycle and suggests that during exit from pluripotency, ESCs were already acquired a propensity towards epiblast vs hypoblast which is dictated by their cell cycle states.

### ESRRB is an essential factor for Retinoic Acid based XEN differentiation

To identify potential genes expressed during pluripotency and which promote XEN differentiation in the context of cell cycle, we first profiled bulk mRNA of ESCs in the G1 and G2/M states. Based on three replicates of G1 and G2/M ESCs, we detected ∼2000 genes expressed at higher levels in G2/M and ∼400 genes expressed at higher levels in G1 (Fig. 3A, Supplementary Table 2). The majority of these genes were cell-cycle regulators, and, as a result, only cell-cycle GO terms were significantly enriched. However, small number of the identified genes are involved in regulation of pluripotency and early differentiation. For example, genes expressed at higher levels in ESCs in the G1 state compared to G2/M included *Nanog, Klf4, Otx2, Leafty1*, and *Pax3*, which are central regulators of EpiSCs initiation ^47^. In the group of genes that show higher expression in G2/M cells we did not find any known direct drivers of the XEN lineage. However, we identified two potential candidates: *Sall4* and *Esrrb* (Fig. 3A). *Sall4* is expressed in both ESCs and XEN cells and encodes a transcription factor that regulates expression of key XEN lineage-associated genes such as *Gata4, Gata6, Sox7*, and *Sox17* ^48^. *Esrrb* encodes a pluripotent transcription factor that is involved in regulation of the trophectodermal lineage ^49^ and, together with Oct4, Sox2, and Klf4, in maintaining ESCs by mitotic bookmarking of pluripotent genes ^50^. In addition, Esrrb is associated with Tfe3, a non-cycled factor that regulates the hypoblast/PrE circuitry ^40^, and together with Gata3, Eomes, Tfat2c and Myc induces a XEN-like state during reprogramming ^51^. To test if the difference in *Esrrb* expression levels observed between G1 and G2/M ESCs is kept also at the protein level, we performed Western Blots (WB) for ESRRB in G1 and G2/M ESCs. Based on five biological replicates, we found small but significant higher levels of ESRRB in G2/M ESCs (Fig.S4).

**Figure 3:**
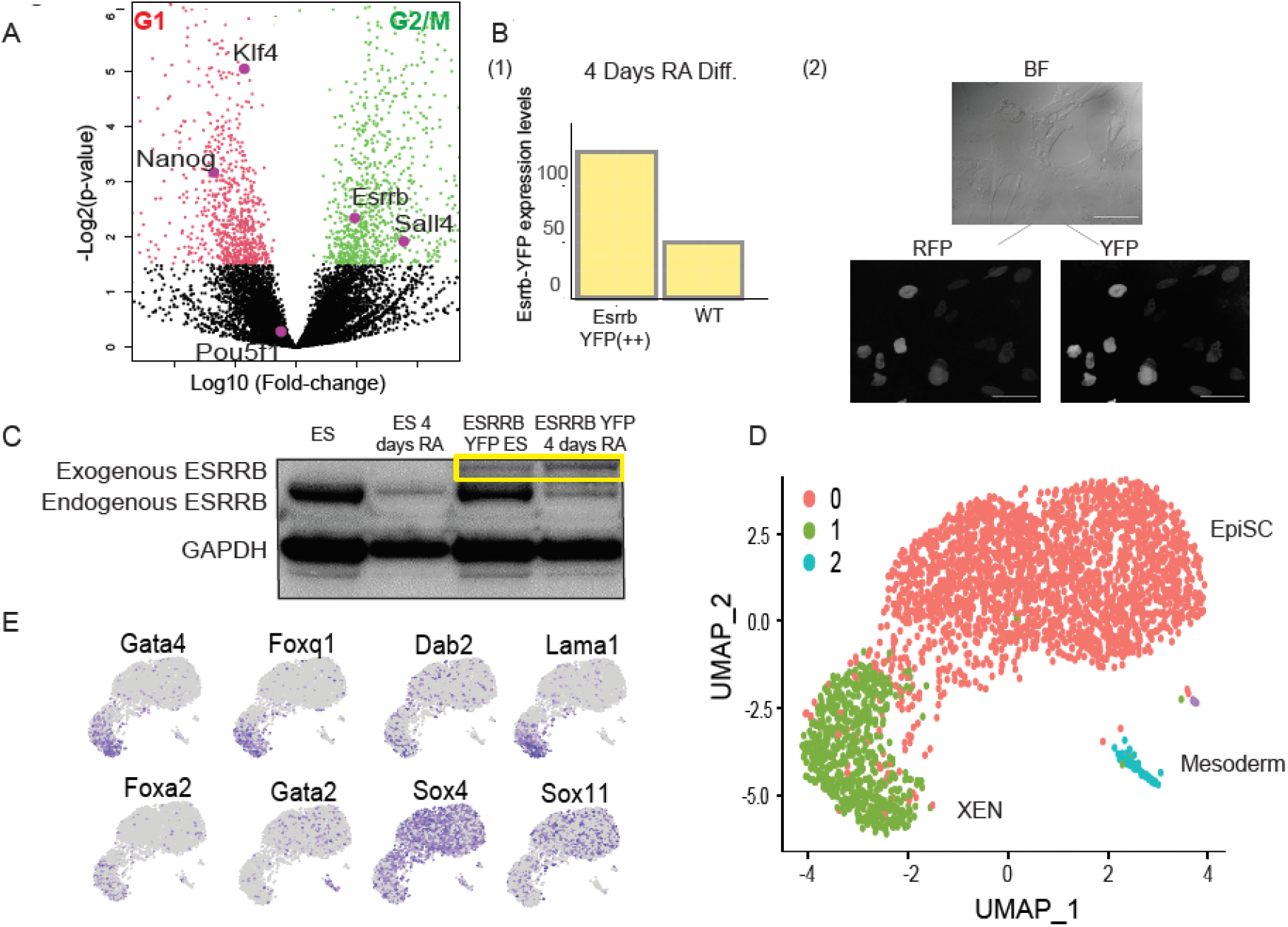
ESRRB is an inducer of XEN cells. (**A**) Bulk RNA-seq data of ESCs in the G1 state (left) vs. the G2/M state (right). The fold change in G2/M vs. G1 ESCs is plotted vs. t-test p-values (n=3). *Esrrb* and *Sall4* genes show G2/M upregulation. *Nanog* and *Klf4* genes show G1 upregulation and *Pou5f1* is insensitive to ESC cell cycle state. (**B**) (1) FUCCI cells non-infected (WT) vs. infected with *Esrrb-YFP* lentiviral construct. Y axis represents expression levels based on RNA-seq read counts of lentiviral transcripts (FUCCI also contributes to the overall count, hence WT counts). (2) Confocal imaging of cells following 4 days with RA showing an overlap between *Esrrb-YFP* and G1 cells (marked by RFP). <u>Upper</u>**-** Representative X20 bright-field image, scale bar 50 µm. <u>Lower</u>-Representative X20 confocal images, scale bar 50 µm. (**C**) Western blot of (left to right): WT ESCs, WT 4 days Diff. with RA, ESCs infected with *Esrrb-YFP*, 4 days RA differentiated cells infected with *Esrrb-YFP*. (**D**) UMAP of 3025 G1 ESCs following 4 days differentiation process clustered into three groups using the Seurat pipeline ^45^. The clusters correspond to EpiSC in red, XENs in green, and mesoderm like in blue. (**E**) UMAP plots highlighting expression levels of indicated marker genes. Gray (low) to purple (high) scale indicates average expression signal.

To further determine and validate Esrrb involvement in G2/M-specific XEN induction, we produced a FUCCI ESC line that overexpresses Esrrb-YFP. We used FACS to select G1 cells that express moderate levels of Esrrb, as indicated by YFP signal. We also imaged these selected cells by confocal microscopy to verify nuclear expression of Esrrb-YFP (Fig. 3B) and validated Esrrb over expression using WB (Fig. 3C). After sorting G1 ESCs using the same parameters as previously described, we differentiated the G1 ESCs for 4 days by treatment with RA and then analyzed 3024 cells from three biological replicates with scRNA-seq. In support with our hypothesis, the subpopulations observed for differentiated G1 Esrrb-YFP ESCs included EpiSC (Fig. 3D) and a small subpopulation of *Gata4* and *Dab2* expressing XEN cells (Fig. 3E). This further points out the involvement of Esrrb in inducing XEN state in a cell cycle dependent manner.

Finally, to directly test the function of ESRRB during the exit from pluripotency, we differentiated ESCs lacking ESRRB for 4 days with RA. The KO was done using CRISPR/CAS9 targeting ESRRB. We validated ESRRB KO using WB (Fig. 4A). Supporting our previous observation, ESRRB KO ESCs contributed only to the formation of EpiSC (Fig. 4B-D). In addition, in line with our previous single cell experiments shown in figure 2, a small population of Gata2 cells (mesodermal like progenitor cells) could also be detected even though unbiased clustering did not highlight it as a standalone cluster.

**Figure 4:**
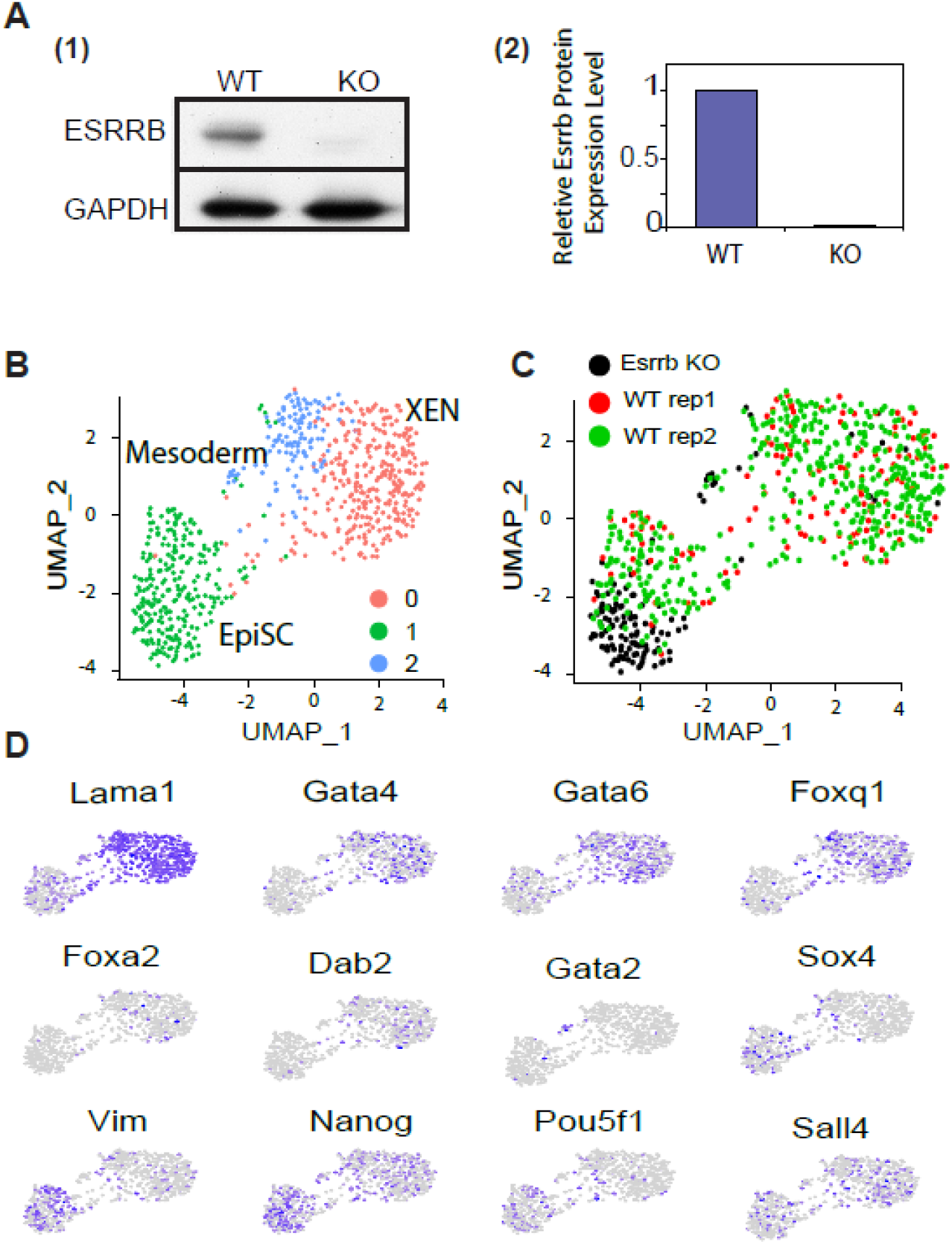
ESRRB-KO ESCs fail to differentiate into XEN cells. (**A**) (1) Western blot of WT and ESRRB-KO ESCs (2) ImageJ pixel based interpartation (y axis: WT based normaliztion) (**B**) UMAP of 700 single cells of 4 days diff. clustered into three groups using the Seurat pipeline ^45^. EpiSC in green, XEN in red and blue. Gata2 Mesodermal like cells are a small subpopulation within the blue cluster. (**C**) The same UMAP as in Fig4B colored for WT (red and green) and ESRRB-KO (black). (**D**) UMAP plots highlighting expression levels of selected marker genes. Gray (low) to purple (high) scale indicates average expression signal.

### During cellular differentiation, linage outcomes are not affected by cell-cycle states

During early differentiation, when progenitor cells emerge, the cells replicate constantly. Therefore, we aimed to explore whether cell-cycle states also influence fate decisions in cells that already exited from their pluripotent state. To this end, we treated ESCs with RA for 2 days, when the main pluripotency factors are significantly downregulated (Fig. S3), and replication time becomes much slower (Fig. 1D). We then sorted the cells based on the cell-cycle phase as described above. Sorted cells were either subjected to scRNA-seq directly or first allowed to differentiate for additional 2 days, followed by scRNA-seq (Fig. S5A). To validate the accuracy of our cell-cycle separation using FACS, we first focused the analysis on previously identified cell-cycle genes ^52^. We calculated the principal component matrix, performed clustering ^53^, and visualized the results using UMAP. The cells clustered based on our FUCCI/Hoechst sorting gates and GO annotations show a very clear separation based on G1 and G2/M states (Fig. S5B). In EpiSC (cluster 0 in Fig. S5C), a separation between G1 and G2/M cells could be detected (Fig. S5D). G1 derived cells expressed higher levels of *Vimentin, Ecm1*, and *Acta2*, ^54,55^ which are associated with later stages of differentiation, whereas G2/M expressed higher levels of *Nanog*, which is associated with premature differentiation status ^56^. Overall, these results suggest that at 2 days following the differentiation signal, the differentiation potential is no longer influenced by the cell cycle state. Interestingly, single cell profiling of cells at day 4 of RA treatment, sorted based on cell-cycle state at day 2 (Fig. S5A), show two major clusters of EpiSC and XEN like cells (Fig. S5B) with similar cell cycle derived proportion of EpiSC and XEN differentiation (Fig. S5C and S5D). Taken together, our study suggests that the differentiation fate of ESCs is strongly influenced by the cell-cycle state at the moment of exposure to the differentiation signal. However, once cells start to differentiate and cellular commitments are made, the cell-cycle becomes irrelevant to fate decisions.

### ESRRB associates with poised enhancers of XEN genes

Esrrb expression is quickly downregulated during the exit from pluripotency. Therefore, to promote XEN differentiation, we hypothesized that in addition to the association of ESRRB with poised enhancers in general, it should specifically mark XEN enhancers. To test this hypothesis, we mapped H3K4me1 and H3K4me2 enhancers of WT and ESRRB KO ESCs. We first extracted H3K4me1 and H3K4me2 signal over TSSs. Surprisingly, we found no significant changes between WT and KO ESCs over promoters and proximal enhancers (Fig.S6). Next we performed an unbiased comparison of peaks over distal enhancers. We extracted differential peaks, hence candidate enhancers, and linked them with their target genes using enhancer atlas ^57^. We found a significant enrichment for WT distal enhancers involved in endodermal differentiation. On the other hand, ESRRB KO ESCs were enriched with enhancers involved in metabolic and apoptotic pathways rather than specific differentiation pathways (Fig. 5A). Having said that, many differentiation genes have several enhancers with additive regulatory effects (Fig. 5B-D). In most cases, ESRRB KO reduced H3K4me1/2 from only one or a small number of enhancers (Fig. 5B) and in many other enhancers, mainly involved in EpiSC and pluripotency, no difference between WT and KO ESCs could be captured (Fig. 5C-D). This might explain why ESRRB KO ESCs can still differentiate to embryonic tissues including endoderm.

**Figure 5:**
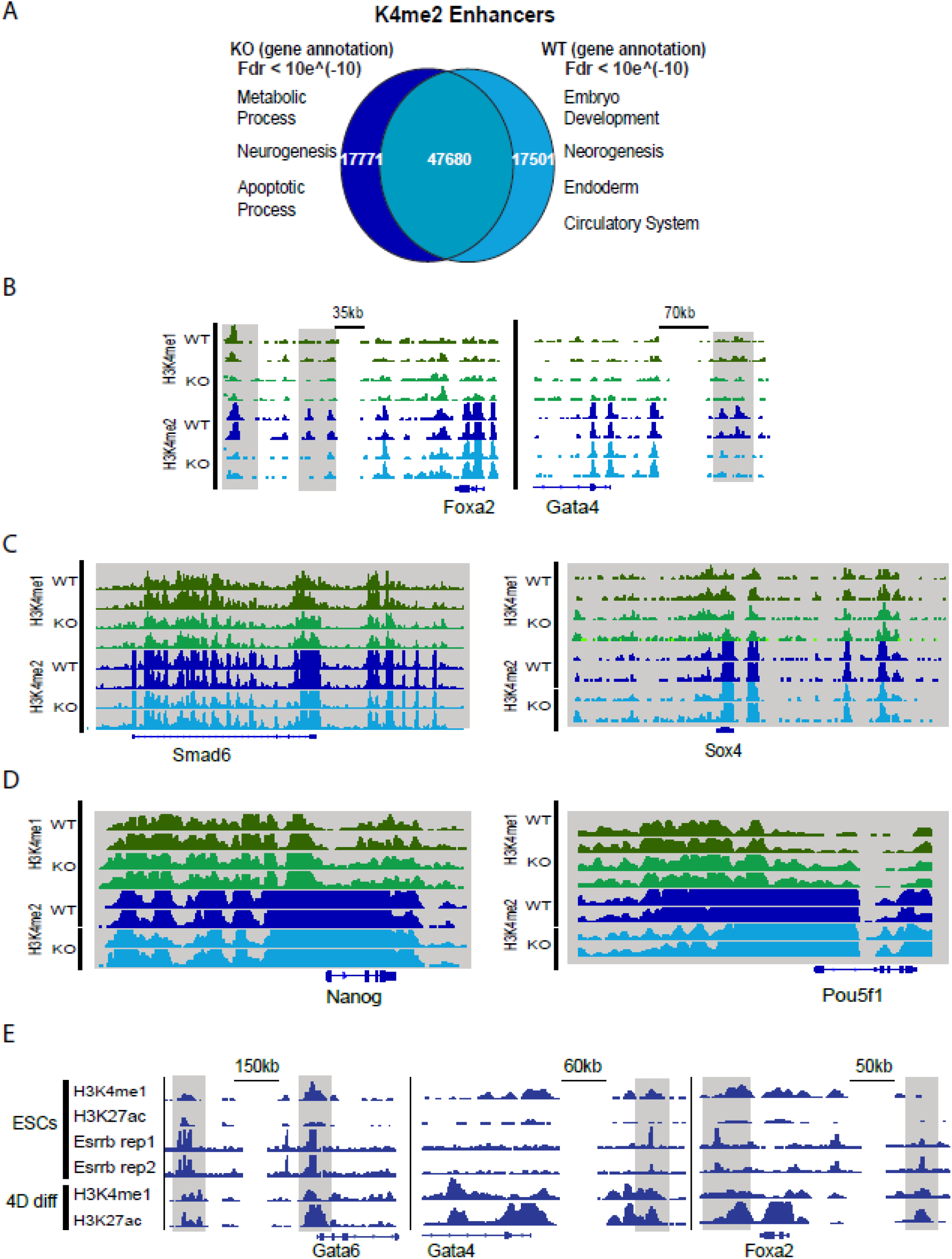
ESRRB associates with poised enhancers of XEN genes. (**A)** H3K4me2 ChIP-seq analysis of distal enhancers comparing WT and Esrrb-KO ESCs. MsigDB enriched annotations correspond to WT vs. Esrrb-KO are shown in each side of the Venn diagram. **(B-D)** ChIP-seq IGV tracks for representative XEN marker genes (*Gata4* and *Foxa2*), EpiSC marker genes (*Smad6* and *Sox4*) and Pluripotent marker genes (*Nanog* and *Pou5f1*) mapped using H3K4me1 and H3K4me2 antibodies for WT and Esrrb-KO ESCs. Distal enhancer regions depleted in Esrrb-KO ESCs marked in grey boxes. (**E)** ChIP-seq IGV tracks of H3K4me1, H3K27ac and ESRRB (Festuccia N. *et al*. ^40^) are shown for ESCs *Gata6, Gata4* and *Foxa2* XEN marker genes. H3K4me1 and H3K27ac are also shown for 4 days RA differentiated cells. Distal enhancers marked with grey boxes.

Next, we mapped H3K4me1 and H3K27ac enhancers in undifferentiated ESCs and in ESCs after 4 days of differentiation. We also obtained and re-analyzed Esrrb ChIPseq maps of interphase ESCs ^41^ enriched for S/G2 ESCs ^11,13^. Both histone marks and Esrrb obtained from ESCs grown in LIF-serum medium condition without 2i. In agreement with our previous findings, examination of XEN marker genes (e.g. Gata6, Gata4, Foxa2, Dab2 and Foxq1) shows that Esrrb was indeed found to be associated with their enhancers. Specifically, these candidate enhancers were poised (H3K4me1 positive and H3K27ac negative) during pluripotency, but were activated upon differentiation as shown by the gaining of H3K27ac (Fig. 5E).

### ESRRB is a XEN specific inducer

To assess if ESRRB is a specific inducer of XEN, we tested two differentiation protocols. We first produced Embryoid Bodies (EBs) ^58^ which contain cells of the three germ layers, Ectoderm, Mesoderm and Endoderm. We observed that EBs size and structure is similar between WT and ESRRB-KO cells (Fig. 6A left panel) and that replication rate is quite comparable (Fig. 6B). Moreover, pluripotent transcription signature of ESRRB-KO ESCs is significantly lower compared to WT ESCs (Fig. 6D1), the overall differentiation scores of EBs ESRRB-KO cells were slightly higher for all differentiation lineages (Fig. S7). We concluded that ESRRB-KO ESCs retain the potential to produce healthy EBs.

**Figure 6:**
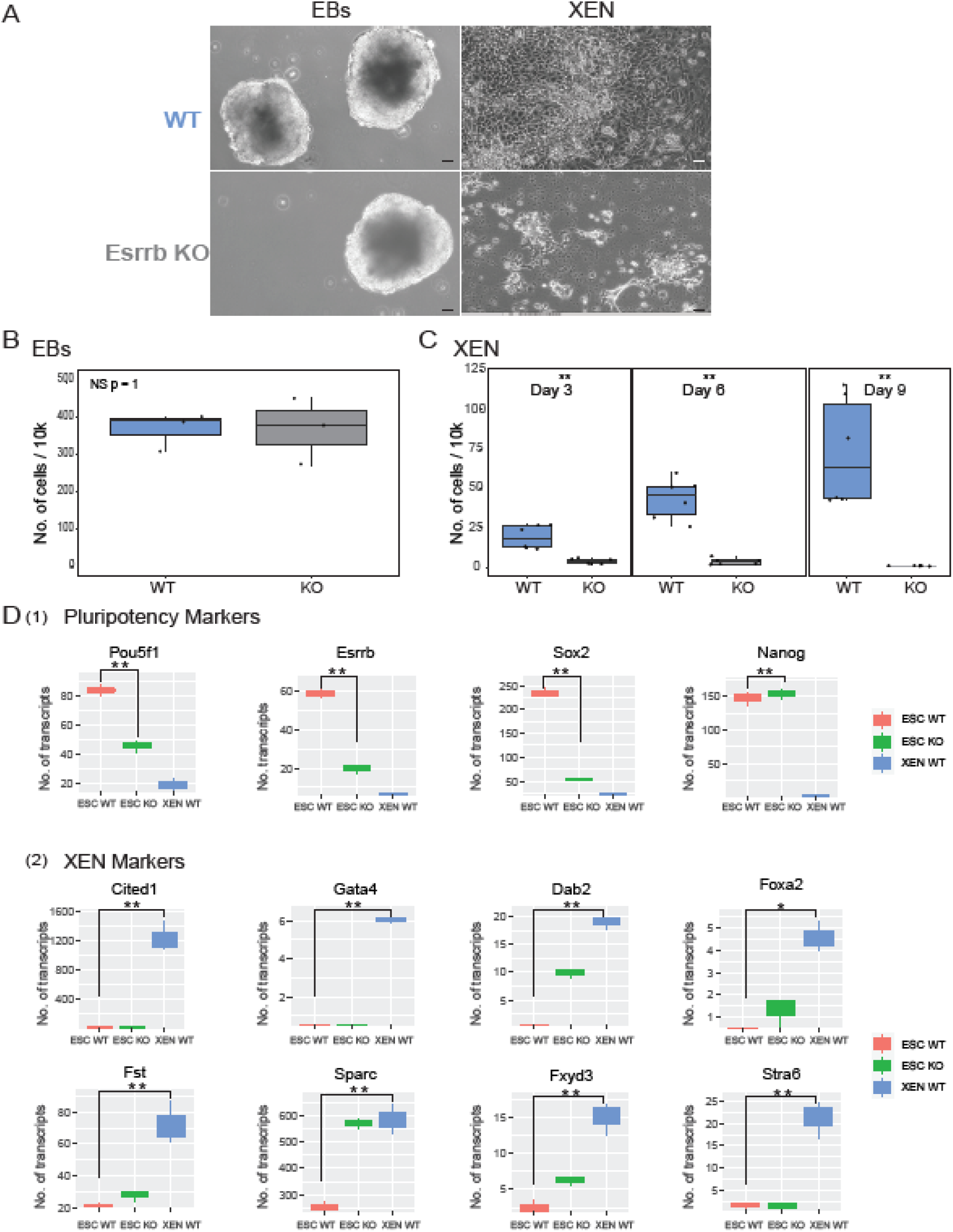
ESRRB is a XEN specific inducer. (**A**) WT and ESRRB-KO ESCs followed by 10 days EBs and 9 days XEN direct differentiation protocols. Representative X10 images, scale bar 50 µm (**B-C**) Live cell count of WT and ESRRB-KO cells for EBs and XEN direct differentiations. Statistics based on three biological replicates (** p < 0.05, * p<0.1 Wilcoxon test) (**D**) Bulk RNA-seq data of WT ESCs, ESRRB-KO ESCs and WT direct XEN differentiated cells (** p < 0.05, * p<0.1 Wilcoxon test). (**1**) Expression levels of selected pluripotent marker genes. (**2**) Expression levels of selected XEN marker genes ^48,60,61^.

Next, we tested 9 days of direct XEN differentiation protocol ^59^. Reassuringly, the majority of ESRRB KO ESCs could not differentiate into XEN but preferred to activate apoptotic pathways (Fig. 6C). Conversely, WT ESCs could efficiently differentiate into cells expressing XEN markers ^48,60,61^ (Fig. 6D2), supporting our initial hypothesis that ESRRB is a specific inducer of XEN.

## Discussion

The molecular mechanisms that specify embryo and extraembryonic germ layer identities are only partially understood. Different transcription factors such as Gata, Fox, and Sox protein families, among others, act to regulate extensive networks in ESCs ^43,56,62^ and are associated with early differentiated states ^63^. However, most of these factors are not expressed during pluripotency, hence they will not play a key role during exit from pluripotency. It has been suggested that cell cycle of ESCs play a key role in specifying differentiation outcomes during exit from pluripotency. To study ESCs cell cycle and the extent in which it dictates differentiation outcomes, we used the FUCCI system ^42^ that allowed us to monitor cell-cycle progression in living cells and combined it for the first time with single cell RNA-seq measurements that allows us to detect the underlying heterogeneity of differentiation. While bulk transcriptome analysis suggested that early differentiation is mainly dictated from G1 ESCs, scRNA-seq in the context of cell cycle identifies heterogeneity in G2/M ESCs differentiation capabilities. These cells included differentiated subpopulations and cells expressing genes associated with a less differentiated state. This suggests that G2/M ESCs contain mixed population of cells states with ESCs predisposed to differentiation with ESCs that are stricter in their pluripotent state while the majority of G1 ESCs are cells prone to differentiation. Therefore, we concluded that population-based measurements make a misleading assumption that only G1 cells are prone to differentiation while averaging G2/M cells hide their underlying heterogeneity.

Our results demonstrate that cell-cycle states of ESCs at the moment they are exposed to RA differentiation signals dictate the decision to differentiate into XENs or EpiSCs/Mesodermal cells. We further demonstrated that Esrrb is a key factor upregulated during G2/M state of ESCs, which promotes XEN differentiation pathway. In agreement with our finding, a previous study showed that Esrrb, in conjunctions with Gata3, Eomes, Tfap2c and Myc, can induce pluripotency by the activation of a unique XEN-like state ^40^. In addition, Esrrb regulates expression of many transcription factors that are critical for maintaining pluripotency and self-renewal ^51^. Betschinger et al. showed that there is significant overlap between chromatin binding of Esrrb and Tfe3, which is a key regulator of the hypoblast/PrE circuitry ^50^. We functionally validated that Esrrb is an inducer of the XEN lineage by: 1. Following overexpression of Esrrb in G1 ESCs, the cells restored the capacity to differentiate into XEN. Thus, the integration of Esrrb into the core transcriptional network of G1 cells stimulated XEN initiation and overcame cell-cycle dependency. 2. We showed that ESRRB KO ESCs lack the potential to form XEN like cells regardless of their cell cycle state at the moment of differentiation activation. This emphasizes that Esrrb is a key regulator of XEN differentiation, and cell cycle dependent expression of Esrrb allow G2/M cells to become XEN like cells. We next validated the importance of Esrrb for XEN formation by applying a direct XEN differentiation protocol. Indeed, the majority of ESRRB-KO ESCs could not produce XEN cells. Reassuringly, EBs formation, a simple differentiation protocol that direct ESCs towards epiblast cells, was not affected by ESRRB-KO.

Other known members of the XEN differentiation circuitry are Tfap2c, Sox17, Eomes, and Cdx2 ^64^. However, we did not detect differential expression of mRNAs encoding these proteins in our scRNA-seq experiments. It is important to note that scRNA-seq suffers from low sensitivity, which predominantly enable the detection of highly expressed genes. In addition, these genes are not expressed in ESCs, which suggests that they are not likely to be involve in the exit from pluripotency step.

Sall4 is critical for XEN differentiation through regulation of expression of *Gata4, Gata6, Sox7*, and *Sox17*, and due to interconnections in the pluripotent regulatory circuitry with Oct4, Sox2, and Nanog ^48^. We observed that *Sall4* expression has a prominent cell-cycle dependency, and its expression was also upregulated during G2/M phase in ESCs. The Sall4 chromatin binding profile is correlated with that of Esrrb, suggesting that Sall4 and Esrrb may be co-regulated during XEN differentiation. Transcription factor recruitment can be modulated by epigenetic modifications to the chromatin ^65,66^ and cell-cycle-specific ESRRB enhancer occupancy could be regulated by methylation of DNA and covalent modification of histone proteins. Further experiments are needed to identify additional epigenetic players that may be involved in the exit from pluripotency specifically to the XEN state.

Overall, the presented results support the tight association between cell-cycle stage and cell-fate determination. We demonstrated that the cell-cycle state affects linage specification only at the exit from pluripotency and although the complete cellular signaling is yet to be comprehensively revealed, Esrrb plays a key role in this regulatory pathway. We therefore suggest a model for which Esrrb accumulation during interphase, from G1 towards G2, expands its enhancer binding capacity towards XEN poised enhancers. Thus, allows the preparation of G2/M specific ESCs towards XEN differentiation (Fig. 7).

**Figure 7:**
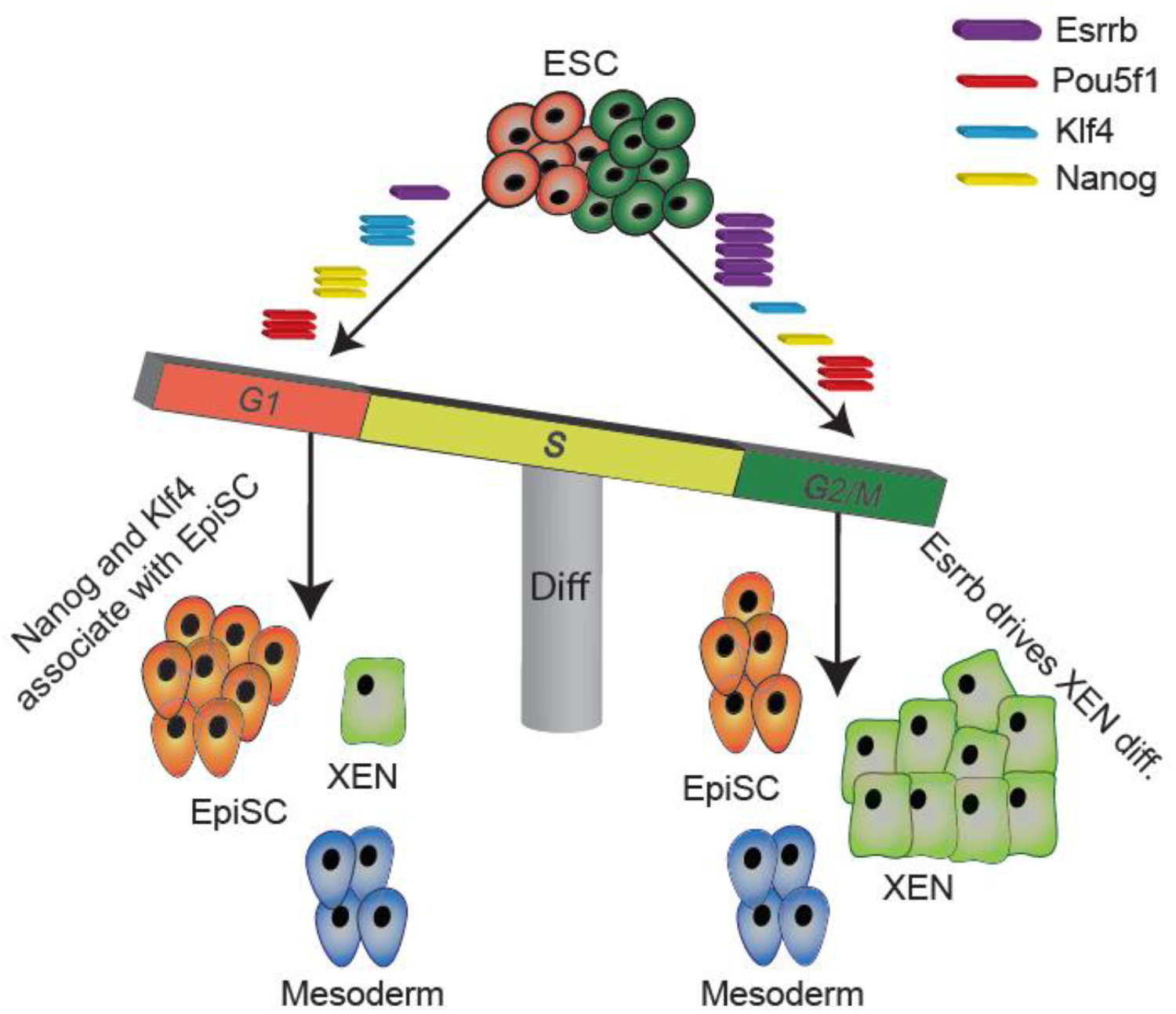
A model for ESRRB dependent XEN induction during exit from pluripotency. ESRRB accumulates in ESCs during interphase with low expression level at G1 phase and high expression during G2/M phase. When cells are at G1, ESRRB’s limited abundance is mainly dedicated for pluripotency maintenance through co-binding with pluripotent factors such as POU5F1, SOX2 and NANOG. When cells are at G2, accumulation of ESRRB allows excess ESRRB to bind XEN specific distal enhancers, thus promoting XEN differentiation.

Further validation experiments and expanding our cell cycle based scRNA-seq protocol on other differentiation pathways such as trophoblast, neuronal and other linage specific differentiations will allow to further increase our understanding of the link between cell cycle and differentiation outcomes exposing other cell cycle or cycling genes that are central for the elusive exit from pluripotency stage and early differentiation decisions.

## Acknowledgments

O.R. is supported by research grants from the European Research Council (ERC, # 715260 SC-EpiCode), the Israeli Center of Research Excellence (I-CORE) program, the Israel Science Foundation (ISF, #1618/16), and Azriely Foundation Scholar Program for Distinguished Junior Faculty. This project has received funding from the European Union’s Horizon 2020 research and innovation programme under the Marie Sklodowska-Curie grant agreement No 765966 - EpiSyStem. We offer special thanks to Prof. Eran Meshorer, Dr. Yonatan Tzur, Prof. Itamar Simon from The Hebrew University for helpful discussions and critical reading of the manuscript.

## Author Contributions

S.H.L., S.F., and O.R. conceived the study and prepared the figures. S.H.L., L.A., S.F., Y.B., and O.R. designed the experiments. S.H.L., L.A., and S.F. M.A. performed tissue culture, FACS sorting, and library preparation for bulk and single-cell RNA-seq. D.B. and X.S. prepared the microfluidics system and performed scRNA-seq experiments. O.R. and S.L. preformed the computational analysis for bulk and single-cell RNA-seq data. S.H.L., L.A., S.F., M.A. and O.R. wrote the manuscript.

## Author Information

All bulk and single-cell RNA-seq data will be deposited in the Gene Expression Omnibus database (GEO). Reviewers, please use this link: in process. The authors declare no competing financial interests. Correspondence and requests for materials should be addressed to O.R.

## Materials and Methods

### Cell culture

Mouse R1 ESCs were a gift from A. Nagy (Lunenfeld-Tanenbaum Research Institute at Mount Sinai Hospital, Toronto, Ontario, Canada). ESRRB Knock out (KO) ESCs shown on Zhbtc4 cell line produced by the Smith lab ^65^. ESCs were seeded on 0.1% gelatin-coated plates (Sigma-Aldrich, G1393) and grown in ES medium (Dulbecco’s modified Eagle’s medium (Sigma-Aldrich, D5671), 10% fetal bovine serum (FBS, Biological Industries, 04-007-1A), 1 mM sodium pyruvate (Biological Industries, 03-042-1B), 0.1 mM nonessential amino acids (Biological Industries, 01-340-1B), 0.1 mM β-mercaptoethanol (Sigma-Aldrich, M3148), 1000 U/ml LIF (Mercury, ESG1107)). The cells were maintained at an incubator at 37 °C with 5% CO_2_ humidified air. For pluripotent conditions, 3 µM CHIR99021 (PeproTech, SM-2520691-B) and 0.2 µM PD0325901 (PeproTech, SM-3911091-B) were added to the ES medium. Cells were passaged every 2 to 3 days using trypsin EDTA (Biological Industries, 03-050-1A). RA differentiation was induced using dissolved RA (Sigma-Aldrich, R2625) in dimethyl sulfoxide (ENCO, 0219605580) at 1 μM in LIF-free media with 10% FBS, on 0.1% gelatin-coated plates. EB’s differentiation was induced using LIF-free media with 10% FBS, on a petri plate.

### Establishment of cell line stably transfected with FUCCI vector plasmids

For cell cycle-phase visualization, the FUCCI (for fluorescent ubiquitination-based cell-cycle indicator) expression system was used ^42^. Plasmids expressing mKO2-hCdt1 (orange-red fluorescent protein) or mAG-hGem (green fluorescent protein) were a kind gift from Prof. Itamar Simon (The Hebrew University, Jerusalem, Israel). 293T cells were kindly gifted from Prof. Meshorer (The Hebrew University, Jerusalem, Israel). Plasmids were transfected into the 293T cells using TransIT-LT1 Transfection Reagent (Mirusbio, MC-MIR-2300) at a ratio of 1:7:3.5 of PMD2.G to psPAX.2 to FUCCI plasmids. The medium was changed after 4 hours to ES medium. After 24 hours, the samples were filtered through a 0.45-mm filter, and cells were resuspended in polybrene-supplemented medium (8 µg/ml PB (Sigma-Aldrich, 107689) in ES medium). After 24 hours, the medium was again replaced with PB-supplemented medium. The cells were then subjected to a two-step FACS sorting. In the first step S, G2, and M cells were sorted by green fluorescence (488 filter) using a FACSAria II cell sorter (Becton Dickinson). The sorted cells were then reseeded and after a week were sorted for red fluorescence (461 filter). For clonal selection, colonies were trypsinzed and seeded in 96-well plates at a density of 1 cell per well.

### Establishment of cell line stably transfected with Esrrb expression vector

Plasmid for expression of Esrrb fused to YFP was a kind gift from Prof. Yosef Buganim (The Hebrew University, Jerusalem, Israel). Plasmids were transfected into 293T p.3 cells using TransIT-LT1 Transfection Reagent (Mirusbio, MC-MIR-2300) at a ratio of 1:7:3.5 of PMD2.G to psPAX.2 to Esrrb-expression plasmid. Medium was changed after 4 hours to ES medium. After 24 hours, samples were filtered through a 0.45-mm filter, and cells were resuspended in PB-supplemented medium. This step was repeated after 24 and 72 hours.

### Fluorescence-activated cell sorting (FACS)

For cell-cycle analysis, cells were trypsinzed and washed twice in PBS. Next, cells were stained with solution containing Hoechst dye (Sigma-Aldrich, B2261). A solution of 9 µl of 25 mg/ml stock of dye in 1 ml of PBS with 2% FBS was used for 5×10^6^ to 6×10^6^ cells. The G1 and S/G2 population were sorted using FACSAria II cell sorter (Becton Dickinson) in PBS supplemented with RNase inhibitor (NEB, M0314L). Sorted samples were processed for scRNA-seq or cDNA-seq.

### Quantification of cellular proliferation rates

To assess cellular proliferation, ESCs cultured with RA were stained with CellTrace Violet Cell Proliferation Kit (Rhenium, C34571) following the manufacturer’s instructions for labeling of adherent cells. Stained cells were analyzed using FACS and by confocal imaging.

### mRNA extraction

mRNA extraction was performed using Invitrogen Dynabeads mRNA DIRECT Purification Kit (61011) according to the protocol included with the kit.

### Library preparation

The library for differential gene expression analysis was prepared using 100-1000 ng of mRNA. First, RNA was fragmented using an RNA fragmentation kit (Ambion, AM8740) following the manufacturer’s protocols. Fragmented RNA was purified with 1.4x reaction volume of AMPure XP beads (Beckman Coulter, A63881) and eluted in 10 μl TE buffer (10 mM Tris–HCl, pH 8.0, 0.1 mM EDTA). The resulting fragmented RNA was reverse transcribed using an oligo dT primer: First, 10 μl of RNA were mixed with 2 μl of 50 μM oligo dT primer (Bio-Rad, 1725038) and 1 μl of 25 mM dNTP mix (NEB, N0447S), and samples were incubated for 3 minutes at 65 °C and transferred to ice. The following components were added to the reaction, for a total volume of 20 μl: 4 μl of 5x First-Strand SuperScript III buffer (Invitrogen, 18080044), 2 μl of 0.1 M DTT (Invitrogen, 18080044), 1 μl murine RNase inhibitor (NEB, M0314L), and 1 μl of 200 U/μL SuperScript III RT enzyme (Invitrogen, 18080044). The reaction product (in the form of a cDNA:RNA hybrid) was purified with 1.5x reaction volume of AMPure XP beads (Beckman Coulter, A63881) and eluted in 16 μl TE buffer as described above. For second-strand synthesis, 16 μl of the digestion reaction product were combined with 2 μl second-strand synthesis (SSS) buffer and 1 μl of SSS enzyme mix of mRNA Second Strand Synthesis Module kit (NEBNext, E6111S) and incubated at 16 °C for 2.5 hours, followed by 20 minutes at 65 °C. The resulting product was purified with 1.5x reaction volume of AMPure XP beads (Beckman Coulter, A63881) and eluted with 20.4 μl TE buffer. cDNA edges were repaired using End-It DNA End-Repair Kit (Danyel Biotech, ER81050) by combining the 20.4 μl sample with 3 μl 10X End-repair Buffer, 3 μl 2.5 mM dNTPs, 3 μl 10 mM ATP, and 0.6 μl END-IT enzyme mix at room temperature for 45 minutes, followed by purification with 1.5x reaction volume of AMPure XP beads (Beckman Coulter, A63881) and elution with 32 μl TE buffer. Klenow-mediated addition of an adenine to the 3’ end of the DNA fragments was performed using the Klenow fragment (3’→5’ exo-) kit (NEB, M0212L) by combining the 32 μl sample with 5 μl 10X Klenow Buffer NEB 2, 10 μl 1 mM dATP, and 0.6 μl 5 U/μl Klenow (3’-5’ exo-) at 37 °C for 30 minutes, followed by purification with 1.5x reaction volume of AMPure XP beads (Beckman Coulter, A63881) and elution with 8 μl TE buffer. Illumina library adaptors were added to the resulting cDNA fragment using DNA ligase (NEB, M2200S) by incubation with 5 μl 2x Ligase Buffer, 0.5 μl adapter Oligo mix (1:10 in H_2_O), and 0.5 μl 1 U/ml DNA Ligase at room temperature for 15 minutes. The ligated product was purified with 1.5x reaction volume of AMPure XP beads (Beckman Coulter, A63881) and eluted in 11.5 μl TE buffer. The resulting libraries were PCR amplified using standard PE1/PE2 full-length primer mix containing Illumina library indices for multiplexing (sequences are given in Supplementary Table 3). Each PCR reaction contained 11.5 μl post-RT cDNA library, 12.5 μl 2x KAPA HiFi HotStart PCR mix (Zotal, KK-KK2601), and 1 μl of 25 μM 2p Fixed (+barcode) and 2p Fixed primer mix (Supplementary Table 3). Amplified libraries were purified using 0.7x reaction volume of AMPure XP beads (Beckman Coulter, A63881) and eluted in 32 μl TE buffer. A 15-μl aliquot of each resulting library was run in a 2% agarose gel, and size selection for the desired 200-800 bp DNA library fragments was performed using PureLink DNA gel extraction kit (Invitrogen, K210012). Library quality was confirmed by Agilent 2200 TapeStation Nucleic Acids System.

The in vitro transcription sequencing (IVT) library was prepared using 75-100 ng of mRNA, extracted as described above. First-strand synthesis was performed using SuperScript III Reverse Transcriptase (Invitrogen, 18080044) and the T7RTPolyT primer (Supplementary Table 3) at 50 °C for 2 hours followed by 15 minutes at 70 °C and 1 minute at 4 °C. To remove unused primers and primer dimers, the reaction product was combined with 20 μl digestion mix containing 5 μl ExoI (NEB, M0293S), 5 μl HinFI (NEB, R0155S), 5 μl ExoI buffer (NEB, B0293S), 5 μl CutSmart buffer (NEB, B7204S), and 30 μl nuclease-free water and incubated for 1 hour at 37 °C and 10 minutes at 80 °C. The reaction product (in the form of a cDNA:RNA hybrid) was purified with 1.5x reaction volume of AMPure XP beads (Beckman Coulter, A63881) and eluted in 13.5 μl TE buffer. For second-strand synthesis, the 13.5 μl digestion reaction product was combined with 1.5 μl SSS buffer and 1 μl of SSS enzyme mix from the mRNA Second Strand Synthesis Module kit (NEBNext, E6111S) and incubated at 16 °C for 2.5 hours, followed by 20 minutes at 65 °C. For linear amplification by in vitro transcription, 16 μl of SSS reaction products were combined with 24 μl HiScribe T7 High Yield RNA Synthesis Kit (NEB, E2040S) reagent mix containing 4 μl T7 Buffer, 4 μl ATP, 4 μl CTP, 4 μl GTP, 4 μl UTP, and 4 μl T7 enzyme mix. The reaction was incubated at 37 °C for 13 hours, and the resulting RNA was purified with 1.3x reaction volume of AMPure XP beads (Beckman Coulter, A63881) and eluted with 20 μl TE buffer. Aliquots of 9 μl of each sample were frozen for backup at -80 °C, 2 μl of each sample was directly analyzed, and the remaining 9 μl were used in subsequent library preparation steps. RNA was fragmented using the RNA fragmentation kit (Ambion, AM8740) by combining 9 μl sample with 1 μl of RNA fragmentation reagent and incubating at 70 °C for 2 minutes. After transfer to ice, 40 μl fragmentation stop mix containing 5 μl fragmentation stop solution and 35 μl TE buffer was added. Fragmented RNA was purified with 1.4x reaction volume of AMPure XP beads (Beckman Coulter, A63881) and eluted in 10 μl TE buffer. The resulting amplified and fragmented RNA was reverse transcribed using a random hexamer primer (IDT) as follows: First, 10 μl of RNA were mixed with 2 μl of 100 μM PvG748-SBS12-RT random hexamer primer (Supplementary Table 3) and 1 μl of 10 mM dNTP mix (NEB, N0447S), incubated for 3 minutes at 65 °C and transferred to ice. The following components were then added to the reaction for a total volume of 20 μl: 4 μl of 5x First-Strand SuperScript III buffer (Invitrogen, 18080044), 1 μl 0.1 M DTT (Invitrogen, 18080044), 1 μl murine RNase inhibitor (NEB, M0314L), and 1 μl of 200 U/µL SuperScript III RT enzyme (Invitrogen, 18080044). Following reverse transcription, the reaction volume was raised to 50 μl by adding 30 μl nuclease-free water, and the resulting cDNA was purified with 1.2x reaction volume of AMPure XP beads (Beckman Coulter, A63881) and eluted in 11.5 μl TE buffer. Library amplification was performed using 12.5 µl 2x KAPA HiFi HotStart ReadyMix PCR kit (Zotal, KK-KK2601) and 1 μl of 25 μM 2p fixed primers (Supplementary Table 3). Amplified libraries were purified using 0.7x reaction volume of AMPure XP beads (Beckman Coulter, A63881) and eluted in 30 μl nuclease-free water. Aliquots of 15 μl of each library were run in 2% agarose gels, and size selection for the desired 200-600 bp DNA library fragments was performed using PureLink DNA gel extraction kit (Invitrogen, K210012). Library quality was confirmed using the Agilent 2200 TapeStation Nucleic Acids System (Agilent).

For scRNA-seq, encapsulation of cells with Reverse Transcription mix (IGEPAL CA-630-Sigma-Aldrich I8896-50ML, SuperScript III Reverse Transcriptase Invitrogen 18080044, Deoxynucleotide (dNTP) Solution Set Ornat N0446S, D,L-DITHIOTHREITOL Bio-Lab 000448235200, TRIS-HCL 1 M STOCK SOLUTIONS PH 7.5 B 1L Sigma-Aldrich T2319-1L, RNase Inhibitor, Murine NEB M0314L) and gels with unique molecular identifiers was performed using the FLUIGENT Smart Microfluidics Pump System with the following flow rate parameters: Cells 100 μL/hour, RT mix 100 μL/hour, BHMs [barcoding hydrogels] 10-20 μL/hour, Oil flow 80 μL/hour. The 4-nL drops were released at a frequency of 15 droplets per second. To release photocleavable barcoding primers from the barcoding beads, the collection tubes were exposed to 6.5 J/cm^2^ of 365-nm light for 10 minutes. Next, to the collection tubes containing the UV-exposed emulsion was added SuperScript III Reverse Transcriptase (Invitrogen, 18080044), and samples were incubated at 50 °C for 2 hours followed by 15 minutes at 70 °C and 1 minute at 4 °C. Each sample was then demulsified by addition of 50 μl perfluoro-1-octanol (Sigma-Aldrich 370533-25G) to release the barcoded cDNA from the droplets. The aqueous phase containing the barcoded cDNA (∼50 μl) was processed for IVT library preparation as described above.

### ChIP

ChIP Antibodies (H3K4me1 abcam ab8895, H3K27Ac abcam ab4729, H3K36me abcam ab9050, 2 µg antibody per 2×107 cells) were incubated with 25 µL protein A Dynabeads (Thermo Scientific) previously washed twice with blocking buffer (0.5% BSA, 0.5% Tween-20 in PBS) for 2 h at 4 °C in blocking buffer. The conjugated beads were washed twice with blocking buffer before adding the lysate. Lysates were prepared as follows: Cells were trypsinized, washed with PBS, and resuspended with 500 μL PBS per 2×107 cells. The same volume of lysis buffer with 100 U/mL MNase was added, and the cells were incubated for 10 min on ice and then for 15 min at 37 °C. Reactions were stopped by adding 20 mM EGTA followed by centrifugation (20,000 g, 2 min). Supernatants were added to the conjugated beads, and samples were incubated overnight with rotation. Supernatants were removed, and the beads were washed twice with RIPA buffer (10 mM Tris-HCl [pH 8], 140 mM NaCl, 1% Triton X-100, 0.1% sodium deoxycholate, 0.1% SDS, 1 mM EDTA), twice with RIPA buffer high salt (10 mM Tris-HCl [pH 8], 360 mM NaCl, 1% Triton X-100, 0.1% sodium deoxycholate, 0.1% SDS, 1 mM EDTA), twice with LiCl wash buffer (10 mM Tris-HCl [pH 8], 250 mM LiCl, 0.5% sodium deoxycholate, 1 mM EDTA, 0.5% IGEPAL CA-630 [Sigma]), and twice with 10 mM Tris-HCl [pH 8]. The beads were resuspended in 45 μL 10 mM Tris-HCl [pH 8] and 20 µg RNase A. The beads were incubated for 30 min at 37 °C, and 60 µg proteinase K was added. The beads were further incubated for 2 h at 37 °C. Proteinase K was inactivated by heating to 65 °C for 15 min, and 50 μL elution buffer (10 mM Tris-HCl [pH 8], 300 mM NaCl, 1% Triton X-100, 0.1% sodium deoxycholate, 0.1% SDS, 1 mM EDTA) was added, and the samples were incubated in 65 °C for 15 min. DNA was purified from supernatants using Ampure XP beads (Beckman Coulter). DNA was eluted with 22 μL 10 mM Tris-HCl [pH 8].

### ChIP library preparation

DNA samples were converted to blunt-ended, phosphorylated DNA using the End-It DNA End-Repair Kit (Lucigen). DNA was purified using ratio of 1:1.8 sample to AMPure XP beads. Adenosine was added to the 3’ end of the DNA fragments using Klenow (3’-5’ exo-) (New England Biolabs). DNA was purified using ratio of 1:1.8 sample to AMPure XP beads. Ilumina adapters were added by ligation using DNA ligase (New England Biolabs). DNA was purified using ratio of 1:1.2 sample to AMPure XP beads. DNA samples were amplified using KAPA HiFi HotStart (Roche) with 25 μM PE1 and PE2 full-length primers containing Illumina library indices for multiplexing (Table 3). Amplified libraries were purified using 0.7x reaction volume of AMPure XP beads and eluted in 22 μL 10 mM Tris-HCl [pH 8]. Aliquots of 15 μL of resulting libraries were run in a 2% agarose gel, and the desired 200-600 bp DNA library fragments were selected and isolated using the PureLink DNA gel extraction kit (Invitrogen). Library quality was confirmed using the Agilent 2200 TapeStation nucleic acids system and the Agilent High Sensitivity D1000 DS DNA kit. The resulting libraries had an average size of 350-550 bp. Size-selected libraries were diluted to 4 nM concentration and combined for paired-end, single index sequencing on the Illumina NextSeq 550 instrument using an illumina 550 High Output v2 (75 cycles) kit. Cycle distribution was 45 cycles for Read 1, 35 cycles for Read 2, and 8 cycles for the library index read.

**Table 3.**
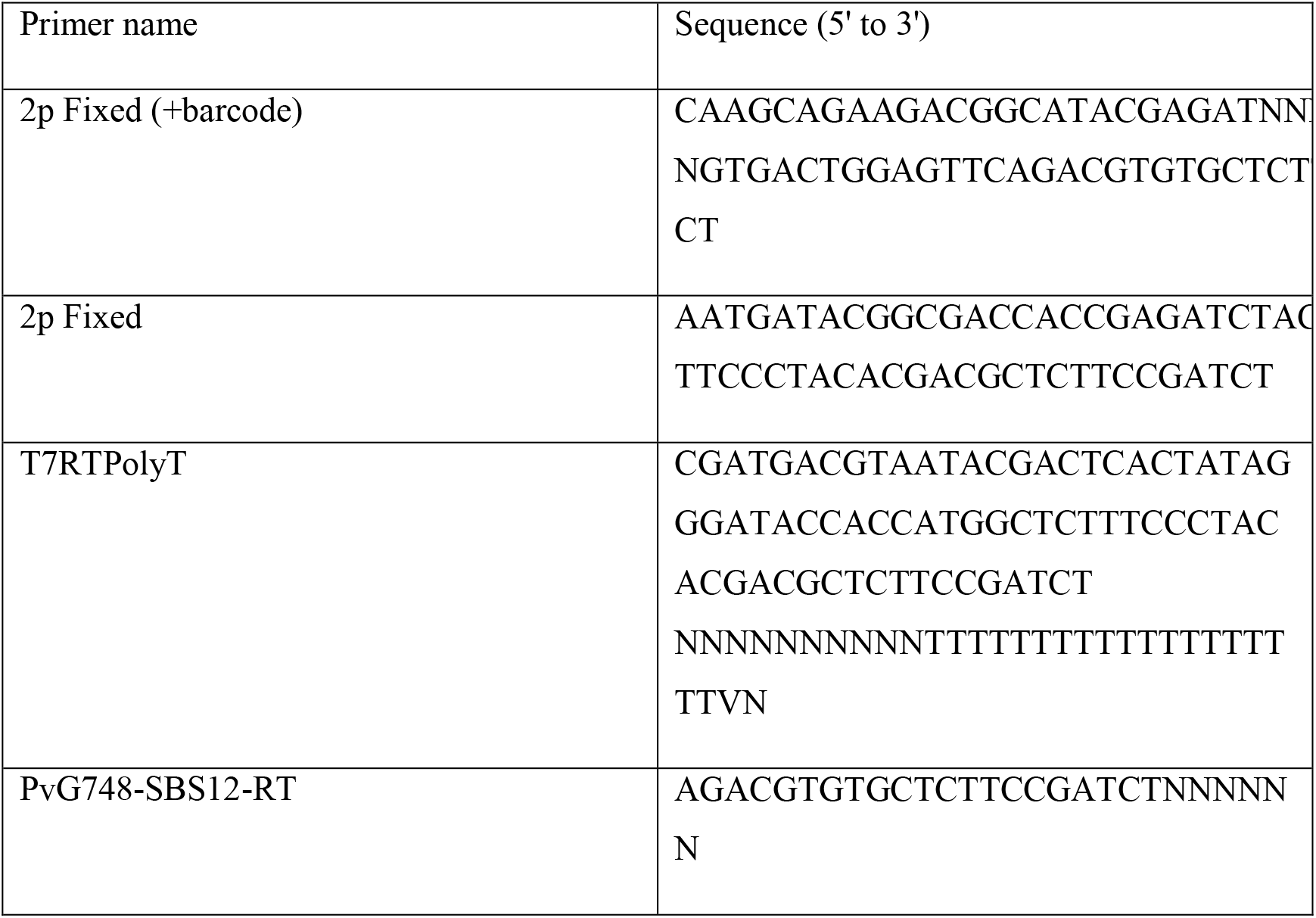
Primer sequences.

**Table 4.**
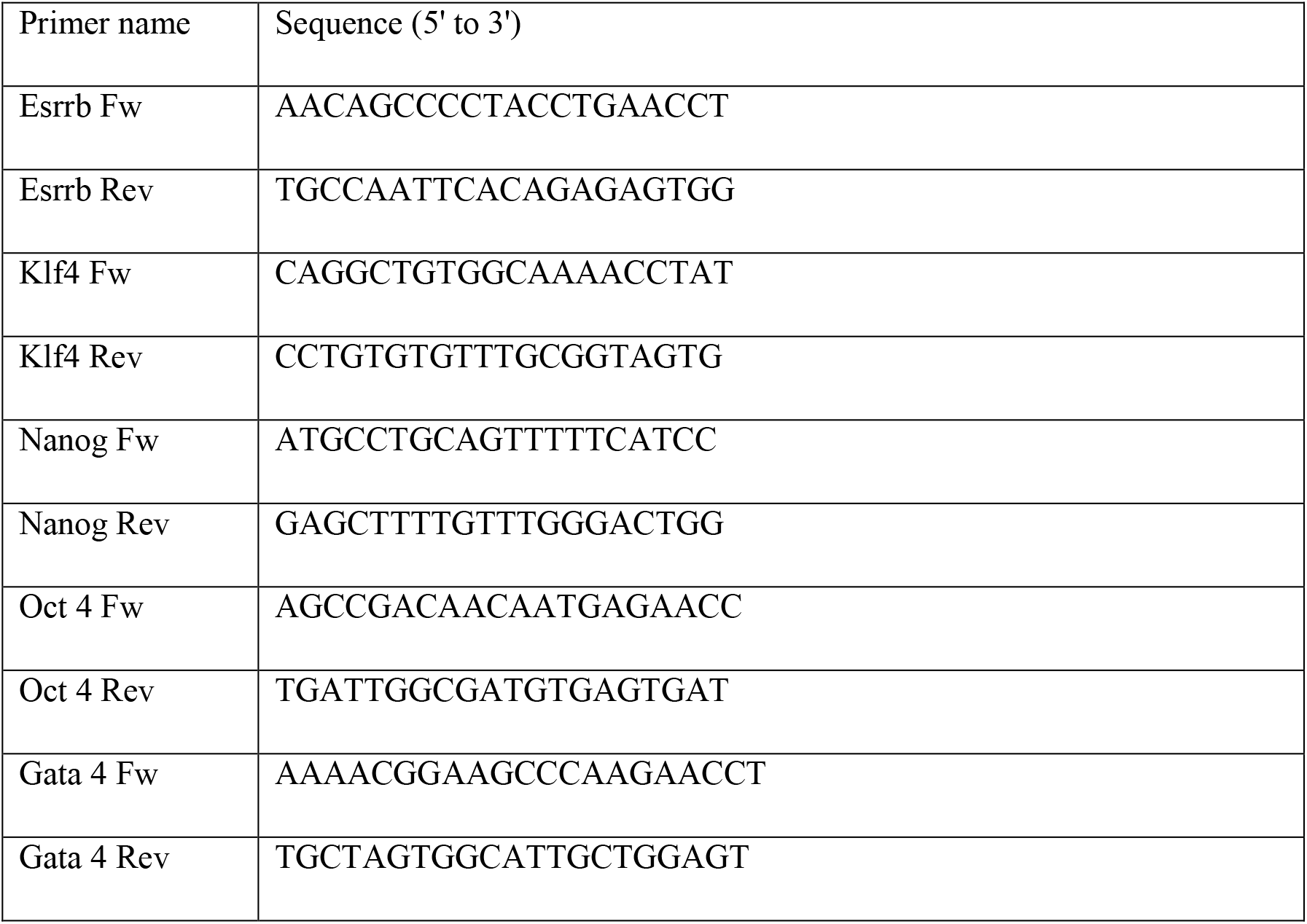
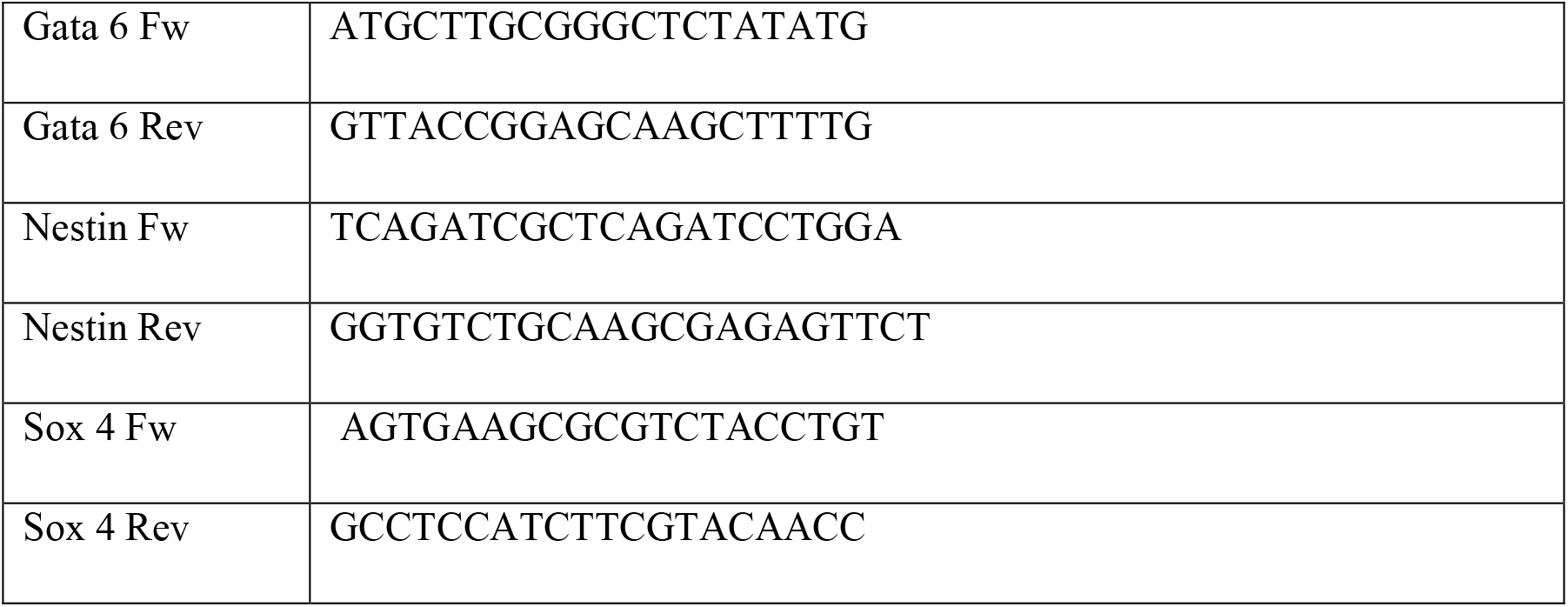
Primer sequences for Real Time PCR.

### ChIP-seq data analysis

Sequencing data were aligned using Bismark and Bowtie (https://www.bioinformatics.babraham.ac.uk/projects/bismark/Bismark_User_Guide.pdf) using paired-ended approach. TDF genomic browser files were produced using IGV count. We applied HOMER to find peaks using ChIP-seq criteria and used BEDTools to intersect bins with genomic intervals such as promoters, genes, and predicted enhancers.

### Real Time PCR

RNA template (500 ng) was used for one-step RT-PCR amplification protocol (SYBR Green I RNA amplification kit, Roche, Basel, Switzerland). Primers were designed using the LC Primer/Probe design software (Roche) and ordered from IDT. All primers are given in 5′ to 3′ direction (Supplementary Table 4) Amplification efficiency for each primer pair was calculated and used for relative quantification. Samples were normalized against GAPDH mRNA levels.

### Western Blot and Antibodies

Proteins were separated by SDS-PAGE on 4%–20% polyacrylamide gradient gels and transferred to 0.45-μm nitrocellulose membranes (iBlot2, PVDF, mini Transfer Stacks, Thermo Scientific; IB24002v). The membranes were incubated with the appropriate primary and secondary antibodies and washed with PBS-Tween 20. Horseradish-peroxidase-conjugated secondary antibodies were detected by SuperSignal West Pico Chemiluminescent Substrate (Thermo Scientific; PI-34080). Antibodies used were anti-human ESRRB /NR3B2 (R&D systems; PP-H6705-00), anti-GAPDH (Abcam; AB-ab8245) Goat Anti-Mouse (Jackson ImmunoResearch; 115-035-062).

## Supplementary Tables

**Table 1** - Single Cell Gene expression analysis during differentiation. Excel sheet is attached.

**Table 2** - Bulk Gene expression analysis during differentiation. Excel sheet is attached.

### Deep sequencing

Deep sequencing was carried out on an Illumina NextSeq using commercially available kits from Illumina (Danyel Biotech FC-404-2005) following the manufacturer’s protocols.

### Data analysis

The Illumina output was analyzed using an in-house Perl script that produced a reads matrix that was aligned using RSEM ^67^ with Bowtie ^68^. The resulting matrix was analyzed in R. For bulk data analysis the transcript per million (TPM) values were used to compare between libraries. Differential gene expression was visualized using volcano plots. Statistical analysis was performed for the two replicates using a two-sided t-test, and p values of <0.05 were deemed significant.

scRNA-seq data was analyzed using the Seurat v2.4 pipeline ^45^. First, cells with more than 10,000 unique molecular identifiers were retained for further analysis. A global-scaling normalization was performed on the filtered dataset using “LogNormalize” with a scale factor of 10,000. Identification of highly variable genes was performed with the following parameters: x.low.cutoff = 0.2, x.high.cutoff = 5, y.cutoff = 0.5, and y.high.cutoff = 10. Cell-to-cell variation in gene expression driven by batch, cell alignment rate, and the number of detected molecules were regressed out and a linear transformation was applied. A principal component analysis was performed on the scaled data with 12 principal components. Clustering was done with resolution of 0.6, and tSNE or UMAP was using for visualization.

### Gene Set Enrichment Analysis

Gene Set Enrichment Analysis was done using GSEA software ^69,70^ with an false negative discovery q value <0.01.

### Statistical analyses

|  | G1-phase | G2-phase |
| --- | --- | --- |
| **Specific cluster: X** | No. of G1 cells in X | No. of G2 cells in X |
| **All other clusters** | Total no. of G1 cells - no. of G1 cells in X | Total no. of G2 cells - no. of G2 cells in X |

Chi-square goodness-of-fit test was performed using “chisq.test” function in R, with the following parameters:

For small numbers, Fisher’s exact test was performed instead.

