## Supplemental Figures for "*Esrrb* is a cell cycle dependent XEN priming factor balancing between pluripotency and differentiation"

Figure S1

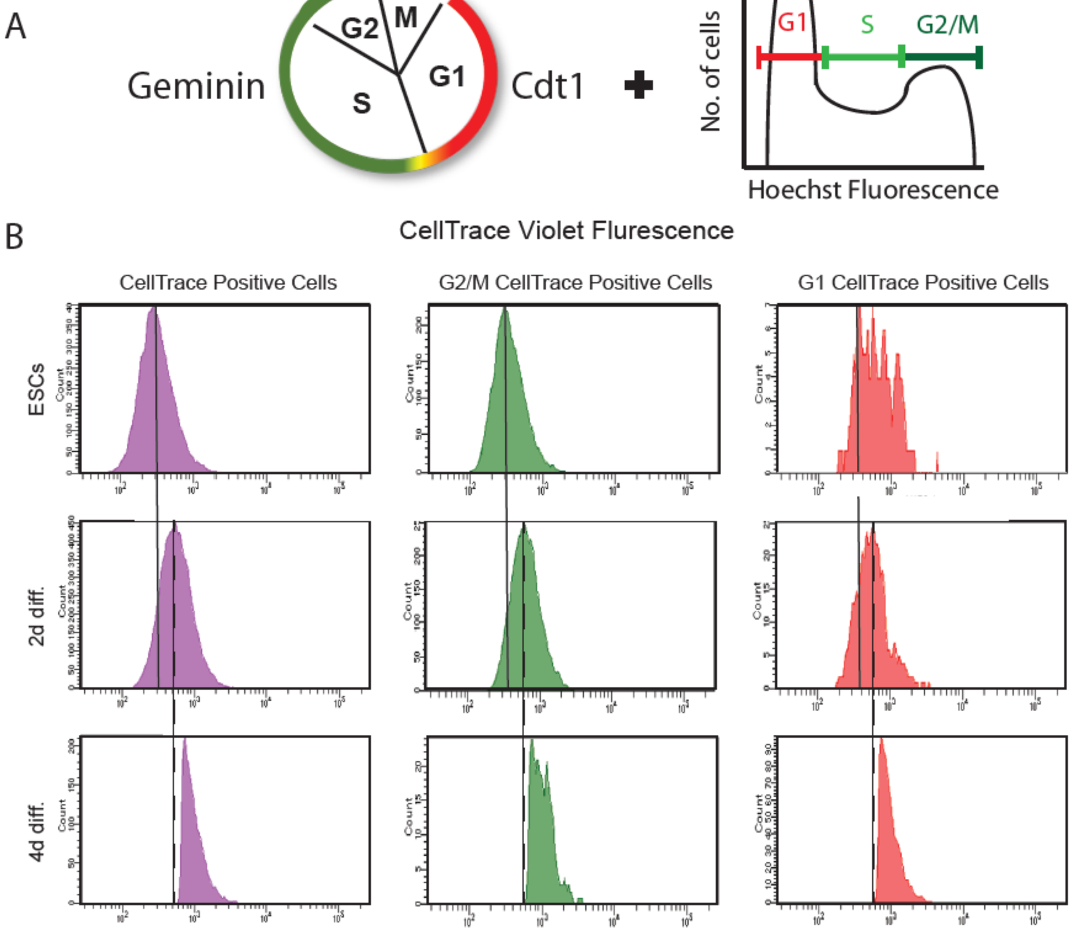


**Figure S1: Cycle-based sorting method**. (**A**) Description of cell cycle-based sorting. Sorting method combined Hoechst staining together with FUCCI expressing cells. (**B**) ESCs were stained with CellTrace Violet (CTV) dye (upper panel) then treated with RA for 2 days (middle panel) and for 4 days (lower panel). Vertical line indicates the mean fluorescence for ESCs. Dashed line indicates mean fluorescence for 2 days diff. cells. Left panel- Unsorted cells. Middle panel- G2/M cells. Right panel- G1 cells.

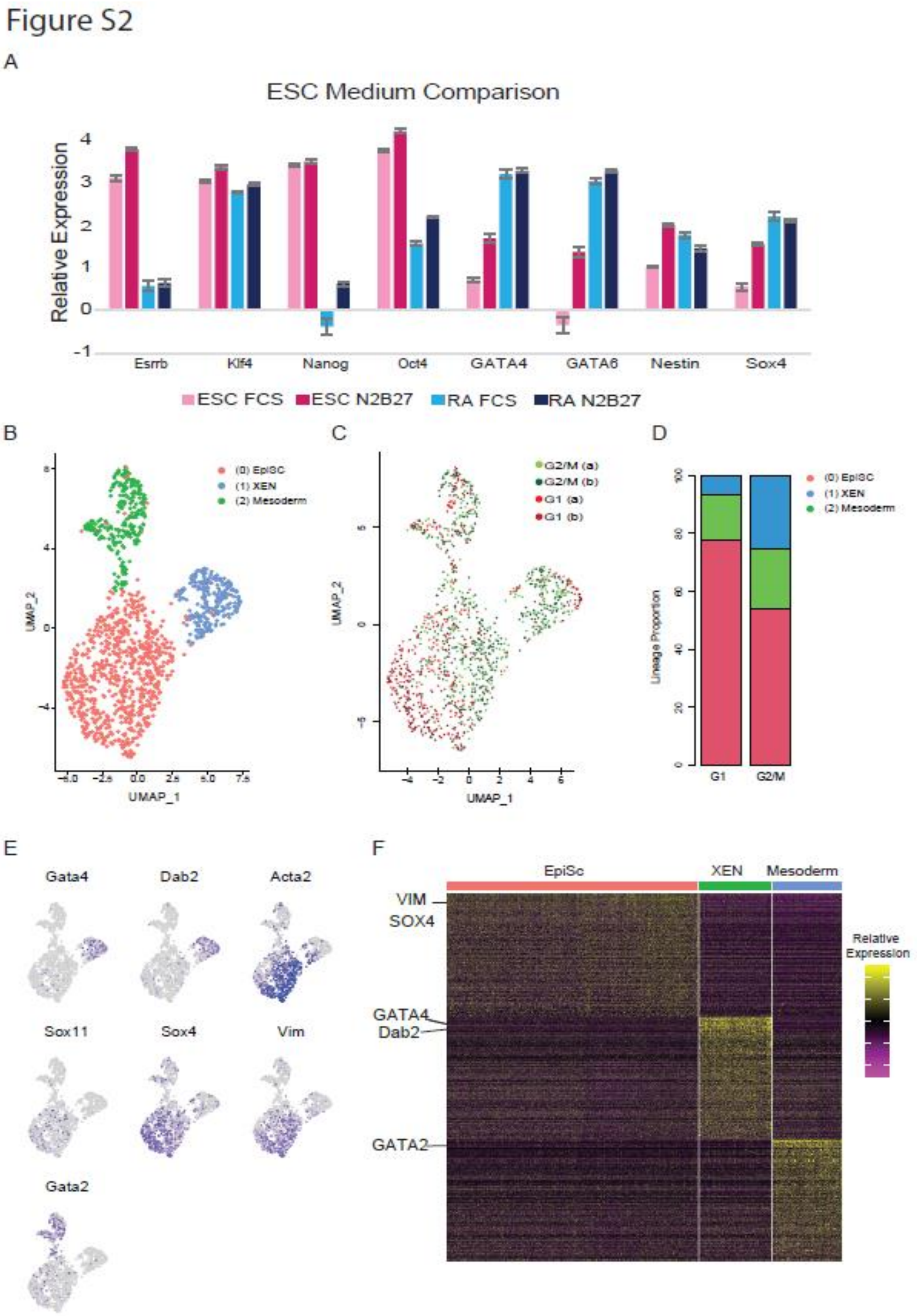


**Figure S2: scRNA-seq analysis of differentiated cells based on G1 and G2/M ESCs populations cultured on chemically defined medium** (**A**) Transcription levels of pluripotent and differentiation markers cultured on FCS and chemically defined medium (N2B27) (**B**) UMAP visualization of ~2000 single cells clustered into four groups using the Seurat pipeline [45]; EpiSC (marked as 0), XEN (marked as 1) and mesodermal-like cluster (marked as 2) are also represented by different colors. (**C**) UMAP visualization of the same single-cell data colored based on the initial sorting of ESCs; G1 cells are in red, and G2/M cells are in green. (**D**) Bar plots representing number of cells in each cluster (**E**) UMAP highlights differentially expressed genes explaining the three different clusters. Gray to purple scale indicates expression levels of each cell. (**F**) Heatmap of gene expression with columns for single cells and rows for genes.

Figure S3

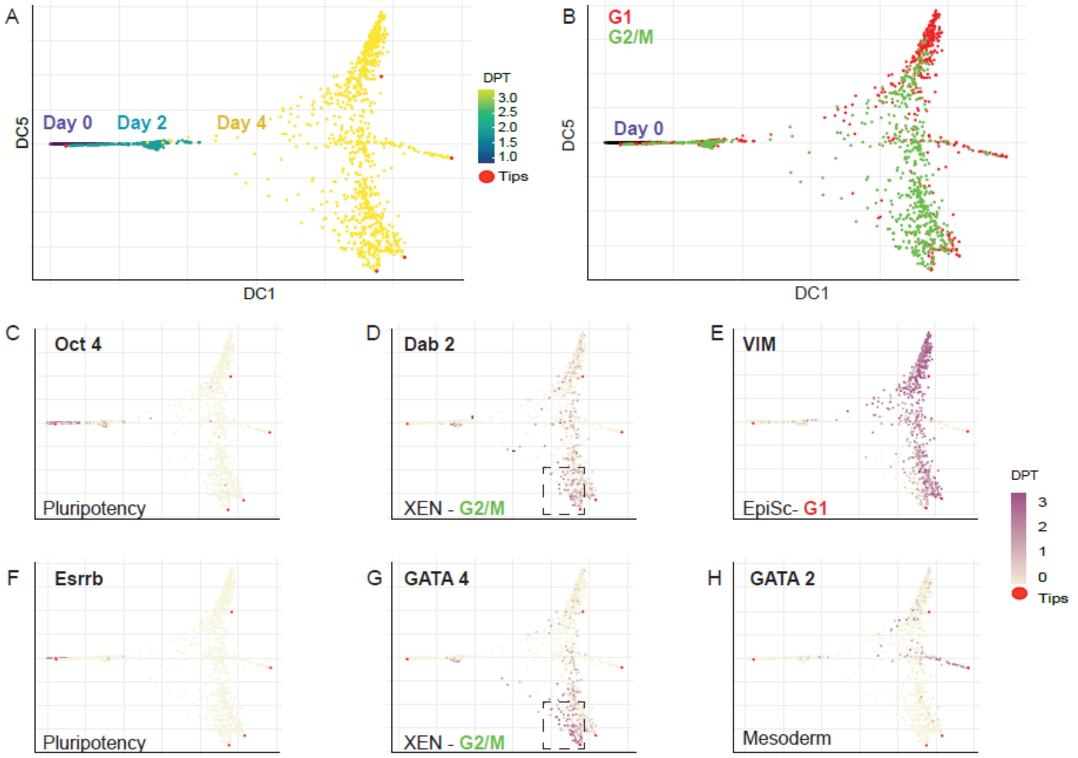


**Figure S3: (A)** Pseudotime visualization of single cell data from 1110 G1 and 1862 G2/M single cells sorted on day 0. ScRNA-seq was done on day 0, 2, 4 of the differentiation processes and clustered using the Seurat pipeline[45]. Clusters correspond to the days of differentiation, respectively. Red dots mark pseudotime roots. **(B)** UMAP visualization of single-cell data colored by G1 cells in red and G2/M cells in green. Cells from day 0 were not colored. **(C-H)** UMAP plots highlighting expression levels of indicated genes. Gray (low) to purple (high) scale indicates average expression signal.

### Figure S4

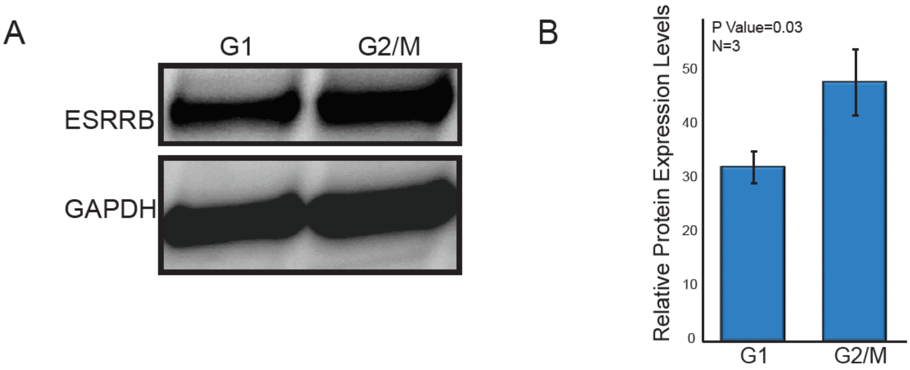


**Figure S4: (A)** Western blot of ESCs after FACS sorting to G1 and G2/M showing small but significant elevated ESRRB protein levels in the G2/M population. **(B)** Relative protein expression levels of ESRRB in G1 cells vs. G2/M cells calculated with the use of Image J software. The scores are the average values of 3 independent experiments.

Figure S5

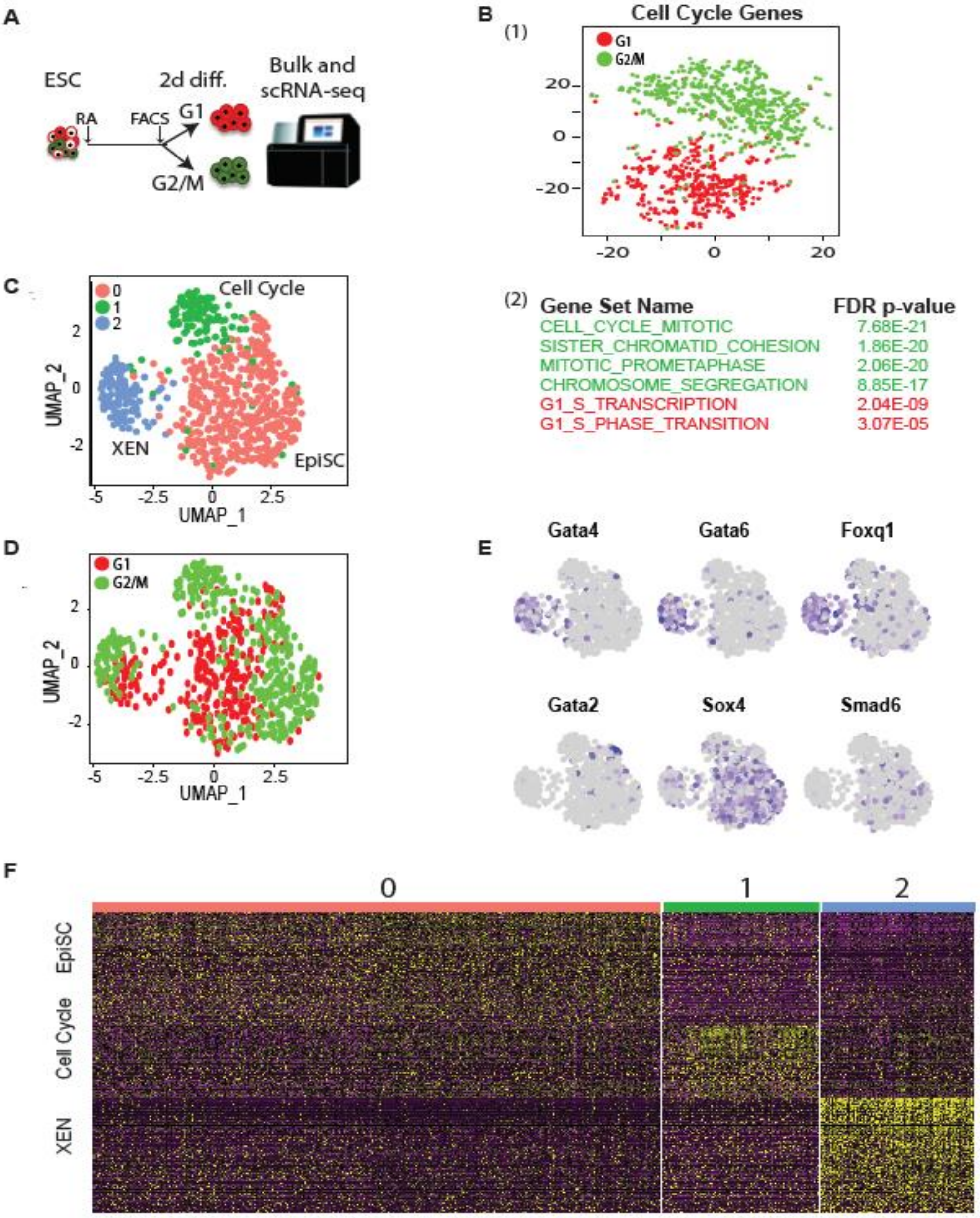


**Figure S5: Bulk and scRNA-seq analysis of sorted G1 and G2/M cells after 2 days with RA**. **(A)** Schematic illustration of the sorting and scRNA-seq experiments: Cells were treated with RA for 2 days and sorted into G2/M and G1 subpopulations prior to bulk RNA-seq. **(B)** (1) tSNE visualization and MsigDB annotations of single-cell data analyzed using only known cell-cycle genes. The two clusters marked in red and green are based on the MsigDB annotations for G1 vs. G2/M cell-cycle states. (2) Gene set names of G1 and G2/M clusters with their FDR p-value **(C)** UMAP visualization of sequencing data from 940 single cells clustered using the Seurat pipeline[45] corresponding to EpiSC in red, XEN cells in blue and cell cycle related subpopulation in green. **(D)** UMAP visualization of single-cell data colored by G1 cells in red and G2/M cells in green. **(E)** UMAP plots highlighting expression levels of indicated genes. Gray (low) to purple (high) scale indicates average expression signal. **(F)** Heatmap of scaled expression based on Seurat normalization and scaling pipeline[43] of the marker genes for each of the clusters.

#### Figure S6

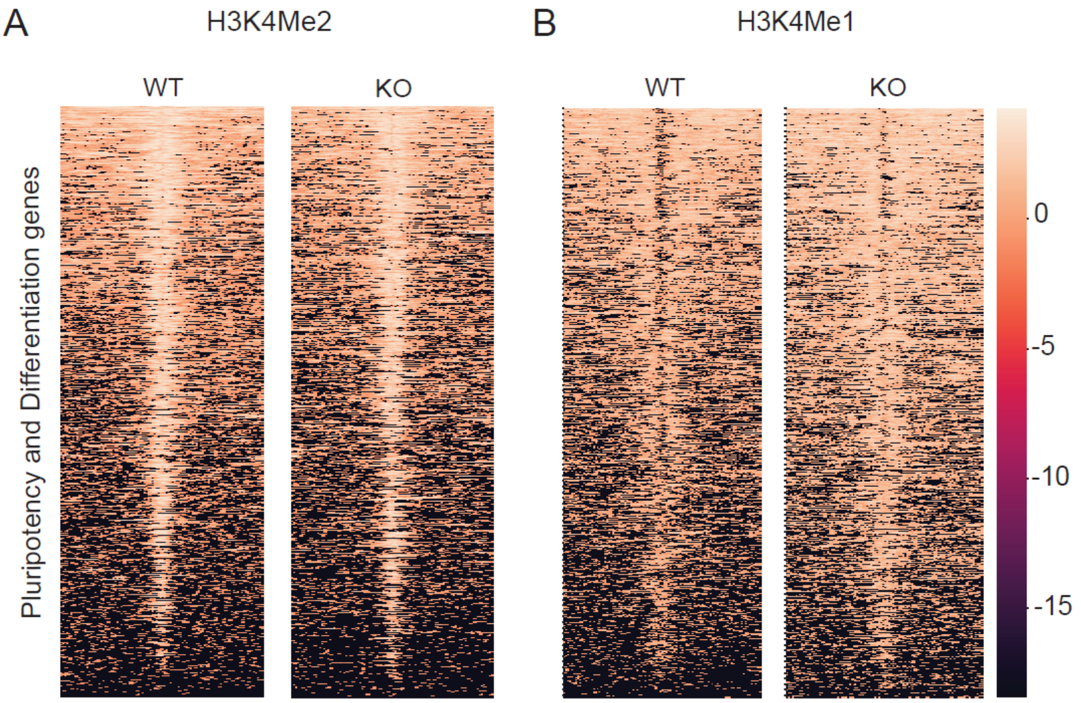


**Figure S6: Heatmap of sequence read density around transcriptional start sites (TSSs), 10 Kb upstream and downstream of TSSs**. (A) H3K4me2 histone marks for WT and ESRRB-KO ESCs (B) H3k4me1 histone marks for WT and ESRRB-KO ESCs.

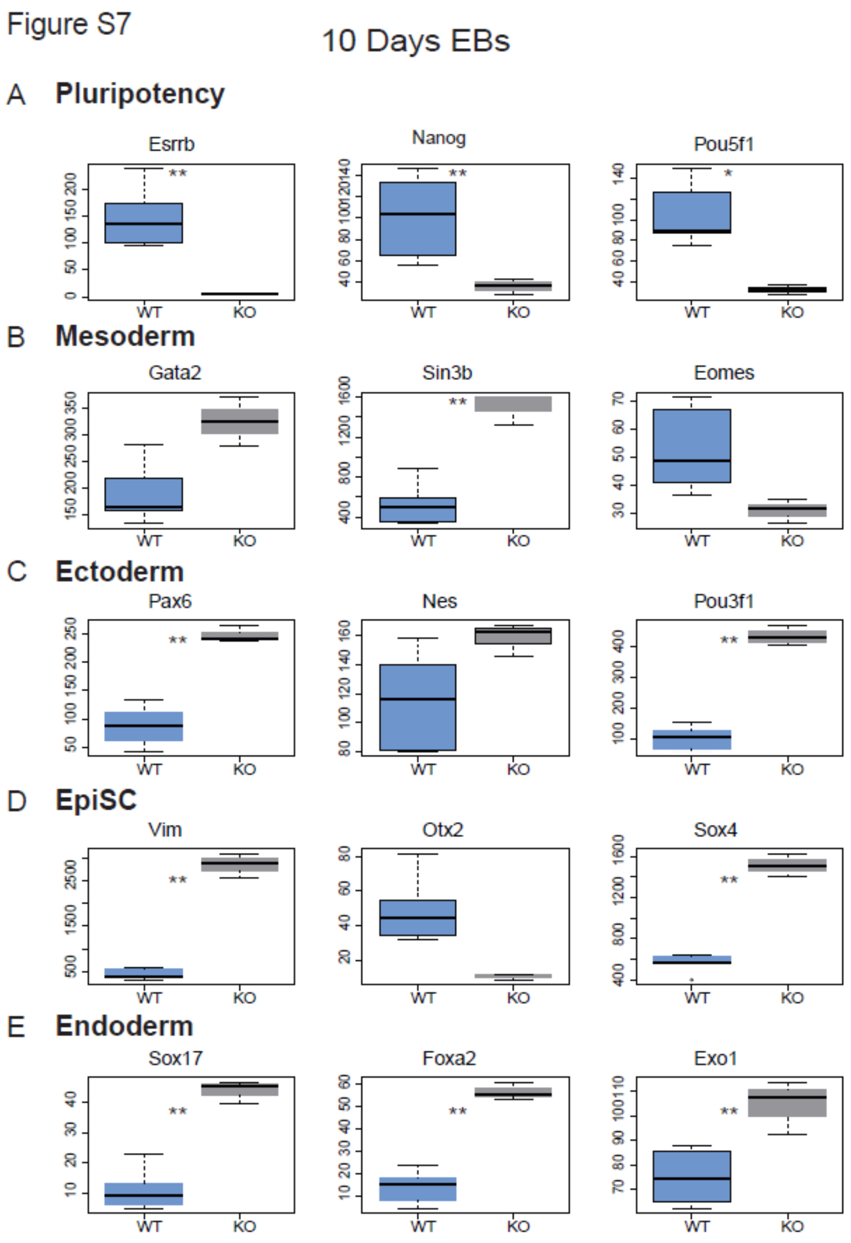


**Figure S7**: **Bulk RNA-seq data of EBs differentiation (10 days).** WT vs. ESRRB-KO cells (**$p<0.05$, *$p<0.1$ Wilcoxon test). Selected marker genes are: **(A)** Pluripotent **(B)** Mesoderm **(C)** Ectoderm **(D)** EpiSC, and **(E)** Endoderm. The overall results indicate ESRRB-KO EBs downregulation of pluripotent genes and upregulation of differentiation genes.
